# The 7SK small nuclear ribonucleoprotein links the cell responses to transcription and replication stress by promoting replication fork reversal and homologous recombination

**DOI:** 10.1101/2025.09.30.678750

**Authors:** Akhil Bowry, Claire Wilson, Richard D.W. Kelly, Rosanna J. Wilkins, Alexandra K. Walker, Christy S. Varghese, Siobhan Murphy-Hollies, Joanna R. Morris, Aditi Kanhere, Eva Petermann

## Abstract

The 7SK-small nuclear ribonucleoprotein complex (7SK-snRNP) plays a crucial role in the response to transcription stress, releasing positive transcription elongation factor P-TEFb to sustain RNA polymerase II activity when transcription is blocked. Many conditions that block transcription also block DNA replication, causing replication stress, and 7SK-snRNP components are putative tumour suppressors with roles in transcription-replication conflicts and double-strand break repair. Here, we investigate potential roles of 7SK-snRNP components in the response to replication stress induced by chemotherapy agents with and without additional impact on transcription. We report that HEXIM1 and LARP7 promote replication fork slowing in response to agents that cause both replication- and transcription stress, in a manner consistent with their canonical 7SK-snRNP functions. Our data suggest that this role in fork slowing is mainly through facilitating RAD51-mediated replication fork reversal rather than transcription-replication conflicts. HEXIM1 and LARP7 promote RAD51 recruitment to replication-associated DSBs or post-replicative gaps under conditions of transcription stress such as induced by camptothecin or BET inhibitors and support HR at direct DSBs. In contrast, HEXIM1 and LARP7 are not required for HR in response to hydroxyurea, which does not cause transcription stress. Our data support that LARP7 roles during replication stress are independent of its reported interaction with BRCA1, and that both LARP7 and HEXIM1 promote survival in response to replication stress-inducing agents. 7SK-snRNP components are not recruited to stressed replication forks and RNA polymerase II inhibition phenocopies loss of these proteins. Taken together, our data support a model where 7SK-snRNP modulation of RNA polymerase II activity helps facilitate RAD51 function under transcription stress conditions, thereby connecting the cell responses to transcription- and replication stress.

## Introduction

Cancer therapies induce cellular stress to kill cancer cells while relying on altered proliferation and stress responses in cancer versus normal cells for specificity. Many cytotoxic cancer therapies damage DNA, causing replication stress and activating DNA repair^1^. DNA damage can also block transcription leading to transcription stress, defined as RNA polymerase II (RNAPII) pausing, stalling or backtracking^2^. It is widely appreciated that replication stress and cellular responses to replication stress play important roles in cancer development and -treatment^3^. How cellular replication stress responses interplay with transcription stress in cancer has received less attention.

Responses to transcription stress include activation of transcription-coupled nucleotide excision repair (TC-NER) and degradation of stalled RNAPII^2^. Another important player in transcription stress is the 7SK-small nuclear ribonucleoprotein (7SK-snRNP), which modulates RNAPII activity in response to a wide variety of treatments that target transcription, including UV radiation and other inducers of bulky DNA adducts, platinum compounds, topoisomerase I inhibitors, and inhibitors of epigenetic regulators such as HDACs, BET proteins and DNA methyltransferases^4–8^.

The 7SK-snRNP is comprised of the non-coding 7SK RNA and the RNA-binding proteins LARP7, MEPCE and HEXIM1^9,10^. The complex binds and inactivates the positive transcription elongation factor b (P-TEFb). Signalling through stress-activated pathways such as through p38 MAPK causes 7SK-snRNP to release active P-TEFb^8,9^ which promotes promoter-proximal pause release of RNAPII by phosphorylating RNAPII and the transcription elongation factors DSIF and NELF^11^ (Figure 1A). Recent work has highlighted that this process is required for a rapid burst in transcription, despite the presence of transcription-blocking DNA damage, of short stress response and non-coding RNA genes that promote survival through largely unknown mechanisms^7,8^.

**Figure 1.**
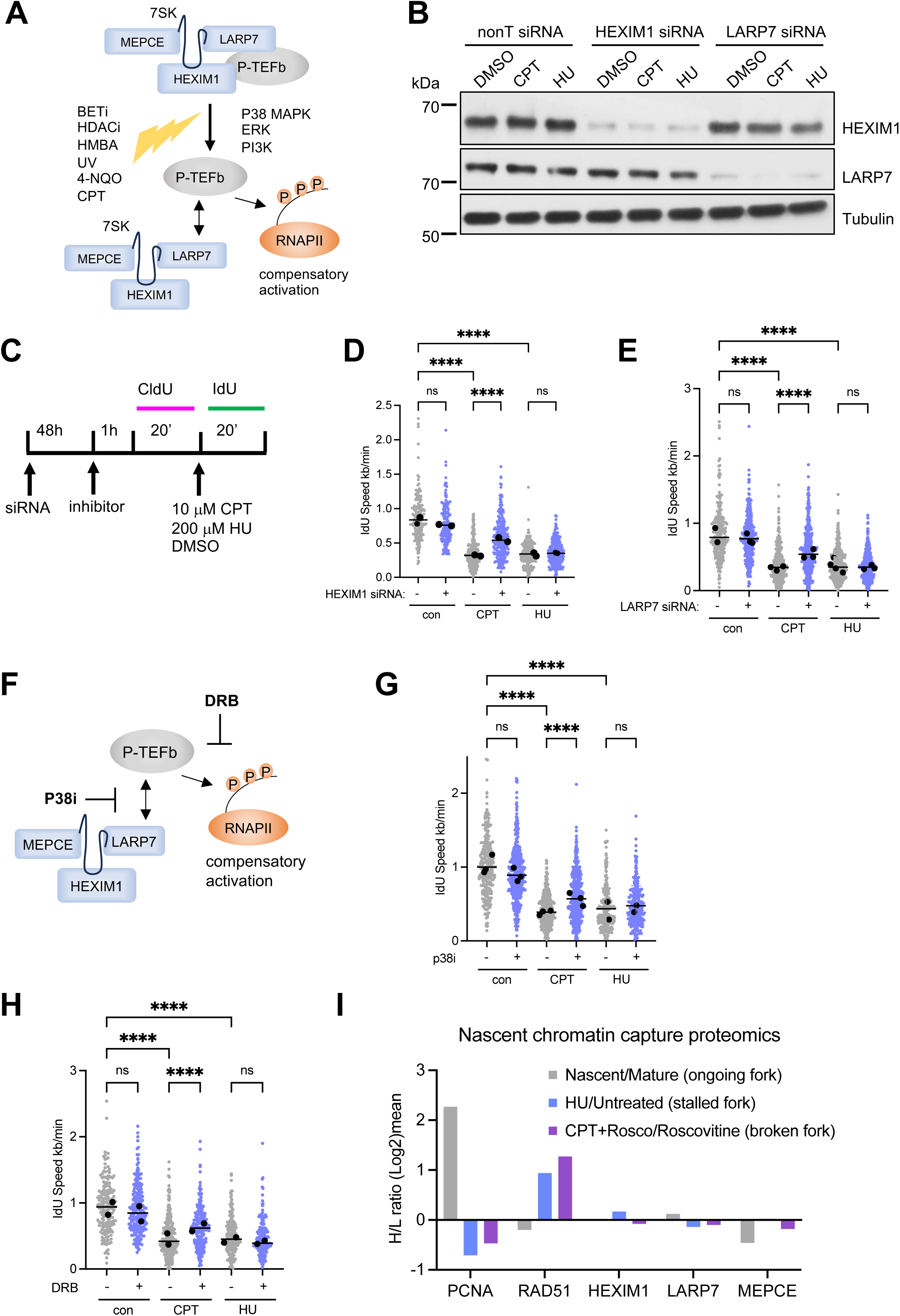
7SK-snRNP components promote fork slowing induced by camptothecin, but not hydroxyurea. A) Schematic of active P-TEFb release from 7SK-snRNP to promote RNA polymerase II (RNAPII) pause release in response to stress and growth stimuli. B) Protein levels of HEXIM1, LARP7 and Tubulin (loading control) in U2OS cells 48 h after siRNA transfection. nonT = non-targeting control siRNA. C) Approach for siRNA depletion or small molecule inhibitor treatment prior to DNA fibre labelling in presence of camptothecin (CPT, 10 μM) or hydroxyurea (HU, 200 μM). D) Replication fork speeds (IdU label) after CPT or HU treatment in presence of HEXIM1 (+) or nonT (-) siRNA. Data from 2 repeats. E) Replication fork speeds (IdU label) after CPT or HU treatment in presence of LARP7 (+) or nonT (-) siRNA. Data from 2-3 repeats. F) DRB inhibits P-TEFb and p38 MAPK inhibitor (P38i) inhibits 7SK-snRNP/P-TEFb dissociation. G) Replication fork speeds (IdU label) after CPT or HU treatment +/- p38i. Data from 3 repeats. H) Replication fork speeds (IdU label) after CPT or HU treatment +/- 100 μM DRB. Data from 2 repeats. I) Nascent chromatin capture proteomics in HeLa S3 cells ^31^ shows limited enrichment or depletion of HEXIM1, LARP7 and MEPCE at ongoing (nascent/mature chromatin), stalled (3 mM HU, 30 min/untreated) or collapsed (1 μM CPT, 40 min + roscovinite/roscovitine) replication forks compared to PCNA or RAD51. Roscovitine was included to prevent new origin firing. Scatter graphs show aggregates and medians (black points) of independent repeats with overall median (line). Kruskal-Wallis with multiple comparisons test, **** p ≤ 0.0001; ns: not significant.

The control of P-TEFb by 7SK-snRNP has been proposed as an anti-cancer mechanism preventing epithelial-mesenchymal transition (EMT) and metastasis^12^. LARP7 and HEXIM1, which are crucial for the function of the 7SK-snRNP, are potential tumour suppressors whose protein levels are decreased in leukaemia and in solid cancers^13^. Such alterations in 7SK-snRNP components could impact the response to transcription blocking cancer treatments.

The transcription-targeting or -blocking agents that induce release of P-TEFb from the 7SK-snRNP frequently also cause replication stress characterised by replication fork slowing. Unresolved replication stress can lead to DNA damage such as single-stranded gaps (ssDNA gaps) or single-ended DSBs (seDSBs), which are implicated in agent toxicity^3^. Homologous recombination (HR) repair, which depends on RAD51, helps repairs ssDNA gaps and seDSBs^14,15^ and is an important determinant of the response to replication stress-inducing cancer therapies^1^. HR factors such as RAD51 also prevent or resolve conflicts between transcription and replication^16,17^, and transcription promotes HR at DSBs^18^. Transcription and replication, as a source of potential conflicts leading to replication stress and DNA damage, are increasingly understood as intertwined in cancer.

In light of this, we previously reported that the 7SK-snRNP component HEXIM1 is required for increased RNA synthesis and RAD51-dependent replication fork slowing in response to the BET inhibitor JQ1^19^. This suggested a role for HEXIM1 in promoting transcription-replication conflicts, but paradoxically, HEXIM1 also prevented DNA damage induction in JQ1-treated cells^19^. Others have since implicated LARP7 or MEPCE in preventing transcription-replication conflicts in absence of BRCA1, and in modulating HR repair at DSBs^20–22^. However, while LARP7 is proposed to prevent HR, MEPCE is proposed to promote HR^20,22^.

Here we aimed to investigate the wider roles of the 7SK-snRNP in the response to replication stress induced by topoisomerase I (TOP1) inhibitor camptothecin (CPT) or ribonucleotide reductase inhibitor hydroxyurea (HU). CPT stabilises TOP1 cleavage complexes, within minutes causing single-strand breaks (SSBs)^23^, torsional stress^24^, RNAPII stalling downstream of RNAPII pause sites, genome-wide changes in RNA:DNA hybrid occupancy, and replication-dependent seDSBs^25,26^. HU causes reactive oxygen species and deoxyribonucleotide triphosphate depletion or imbalance, which both slow and stall replication forks^27,28^.

We report that HEXIM1 and LARP7 promote replication fork slowing in response to agents that cause both replication- and transcription stress, in a manner consistent with their canonical 7SK-snRNP functions. Our data support that this role in fork slowing is mainly through facilitating RAD51-mediated replication fork reversal. HEXIM1 and LARP7 promote RAD51 recruitment to DNA damage sites, HR repair and cell survival especially under conditions of transcription stress such as induced by CPT, suggesting that the 7SK-snRNP links the cell responses to transcription- and replication stress.

## Results

### The 7SK-snRNP promotes replication fork slowing in response to CPT, but not HU

To investigate functions of the 7SK-snRNP in replication stress, we used siRNA to deplete human U2OS osteosarcoma cells of HEXIM1 or LARP7 (Figure 1B), combined with DNA fibre analysis to investigate whether loss of 7SK-snRNP components alters replication fork progression in presence of replication stress-inducing agents. Cells were labelled with CldU to mark ongoing forks, followed by IdU combined with CPT or HU to measure replication fork progression during drug treatment (Figure 1C). The concentrations of both drugs were chosen to reduce replication fork speeds to a similar extent, as higher HU concentrations would block replication fork progression (Figure 1D). As we previously observed for JQ1^19^, HEXIM1 depletion partially rescued replication fork slowing induced by CPT. In contrast, HEXIM1 depletion did not rescue replication fork slowing induced by HU (Figure 1D). While CPT induces many more DSBs than HU, HEXIM1 depletion had also rescued fork slowing induced by JQ1, which does not induce DSBs ^19^. A similar partial rescue of treatment-induced fork slowing was observed when using lower CPT concentrations (Figure S1A), a different cell model, human colon cancer cell line HCT116, and a different drug that releases P-TEFb, HMBA^29^ (Figure S1B), or when LARP7 was depleted (Figure 1E). CldU fork speeds were unaffected by HEXIM1 or LARP7 depletion (Figure S1C, D).

Both HEXIM1 and LARP7 are required for the release of active P-TEFb in response to stress. We therefore investigated whether inhibiting p38 MAPK, which has been implicated in promoting release of P-TEFb from the 7SK-snRNP^8^, or inhibiting CDK9, the catalytic subunit of P-TEFb (Figure 1F), could also rescue CPT-induced fork slowing. Both p38 inhibition with SB203580 and CDK9 inhibition with 5,6-Dichlorobenzimidazole 1-β-D-ribofuranoside (DRB) partially rescued replication fork slowing induced by CPT but did not rescue fork slowing induced by HU (Figure 1G, H). Taken together, these data support that the 7SK-snRNP complex and its release of active P-TEFb promote replication fork slowing in presence of CPT as well as JQ1 and HMBA.

To investigate whether the 7SK-snRNP is recruited to stressed replication forks, we interrogated published replication fork proteome data. Proteomics studies using iPOND^30^ or nascent chromatin capture combined with stable isotope labelling by amino acids in cell culture (SILAC)^31^ (Figure 1I) did not show enrichment of HEXIM1, LARP7 or MEPCE at ongoing, stalled or collapsed replication forks, which suggests that these proteins do not act directly at forks.

### CPT, but not low dose HU, activates a transcription stress response

We next compared the impact of CPT, JQ1 and HU on transcription stress and P-TEFb dynamics. To measure the relocation of P-TEFb, cells were fractionated into a soluble (cytoplasmic and nucleoplasmic) fraction by extracting whole cells with low salt buffer conditions, and a chromatin fraction by extracting chromatin pellets with high salt buffer conditions^32^. P-TEFb moves from the soluble nucleoplasmic into the chromatin fraction in response to CPT, which correlates with P-TEFb release from the 7SK-snRNP^5^. Fractions were probed for CDK9 using Western blot (Figure 2A, B). Quantification of CDK9 signal intensity followed by normalisation to control demonstrated that as expected based on previous reports^4,5^, both JQ1 and CPT treatment increased P-TEFb in the chromatin fraction. In contrast, HU treatment slightly increased the fraction of P-TEFb in the soluble fraction (Figure 2C; Figure S2A). RT-qPCR showed that expression of transcription stress-responsive genes *SERTAD1* and *EGR1*^8,19^ increased in response to CPT, but not HU (Figure 2D).

**Figure 2.**
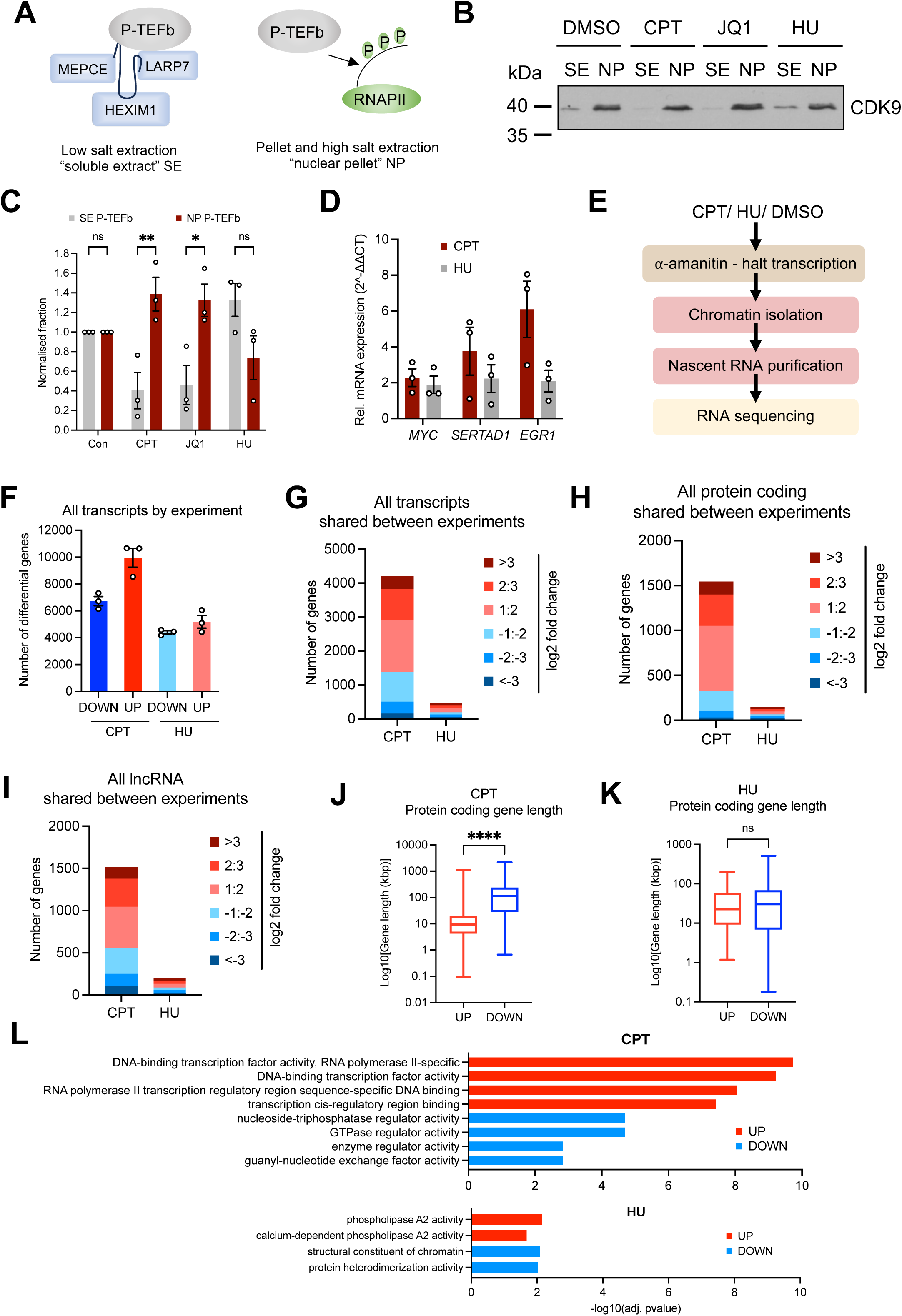
Camptothecin, but not hydroxyurea, activates a transcription stress response. A) Principle of detecting soluble and chromatin-bound P-TEFb by cell fractionation. B) Protein levels of CDK9 in soluble extract (SE) and nuclear pellet (NP) fractions after 2 h treatment with 10 μM CPT, 1 μM BET inhibitor (JQ1) or 200 μM HU. C) CDK9 in SE (7SK-snRNP-bound) and NP (chromatin-bound) P-TEFb calculated as fraction of total CDK9 for each treatment and normalised to untreated controls. n=3. D) RT-qPCR analysis of *MYC*, *SERTAD1* and *EGR1* expression after 2 h treatment with 10 μM CPT or 200 μM HU. n=3. E) Principle of chromatin RNA-seq approach. F) Numbers of nascent transcripts significantly up- or downregulated after 2 h 10 μM CPT or 200 μM HU treatment in each repeat (log_2_-fold change). G) Numbers of up- or downregulated transcripts shared between all repeats. H) Numbers of up- or downregulated protein coding transcripts shared between all repeats. I) Numbers of up- or downregulated lncRNA transcripts shared between all repeats. J) Gene lengths of shared transcripts up- or downregulated in response to CPT. K) Gene lengths of shared transcripts up- or downregulated in response to HU. L) Gene ontology (molecular function) of genes significantly up- or downregulated after CPT or HU treatment. Column graphs show mean ± SEM. 2-way ANOVA or Man-Whitney test, * p ≤ 0.05; ** p ≤ 0.01; **** p ≤ 0.0001; ns: not significant.

To investigate the impact of CPT and HU on transcription stress, we used chromatin RNA-seq to quantify nascent transcripts^33^. Chromatin-bound nascent RNA was isolated 2 h after CPT or HU treatment and subjected to next-generation sequencing (Figure 2E, Figure S2B-D). Data from three biological repeats showed that both CPT and HU treatments altered the synthesis of large numbers of transcripts in every experiment. CPT affected more transcripts than HU, with more transcripts up-regulated than down-regulated (Figure 2F). When we considered only transcripts that were significantly altered across all three biological repeats, this revealed that only CPT induced a specific transcriptional response (Fig. 2G). In addition to protein coding transcripts, transcripts consistently altered in response to CPT included non-coding RNAs such as lncRNAs, miRNAs, and snRNAs (Figure 2H, I, Supplemental Table S1). Nascent RNA sequencing (RNA-seq) had previously shown that both CPT and 4-NQO lead to a rapid (<1 h) relative decrease in longer transcripts and relative increase in shorter transcripts^8,34^. Transcripts up-regulated in response to CPT were also shorter than down-regulated transcripts, consistent with transcription stress (Figure 2J). In contrast, the transcripts consistently up- or downregulated in HU treatment were of similar length (Figure 2K). Protein coding genes up-regulated in response to CPT were enriched in genes involved in the regulation of RNAPII transcription and nucleic acid binding (Figure 2L). These included genes that were previously found up-regulated in response to short (≤ 2 h) treatments with CPT, JQ1 or 4-NQO^8,34,35^ such as the transcription factors *JUN, FOS, EGR1* and *SERTAD1*, *HEXIM1* itself, and ribosomal protein genes. Comparatively fewer gene ontologies were enriched for genes down-regulated by CPT or altered in response to HU. Cell cycle or DNA damage response genes were largely unaffected by either treatment (Supplemental Table S1). Overall, these data are consistent with CPT, but not HU, inducing the P-TEFb relocation and transcriptional changes associated with a transcription stress response. This provides us with tools to investigate the roles of HEXIM1 and LARP7 in presence and absence of overt transcription stress.

### Limited evidence for HEXIM1 promoting transcription-replication conflicts

We speculated that CPT-induced release of P-TEFb might promote conflicts between transcription and replication, as previously suggested for JQ1^19^. We first tested whether HEXIM1 or LARP7 depletion would prevent the re-localisation of P-TEFb to chromatin. HEXIM1 and LARP7 depletion attenuated the CPT-induced increase in chromatin-bound P-TEFb (Figure 3A-D; Figure S3A-D). However, especially in the LARP7 depletion experiments, a significant fraction of CDK9 could still relocate to chromatin (Figure 3B, D, Figure S3C, D).

**Figure 3.**
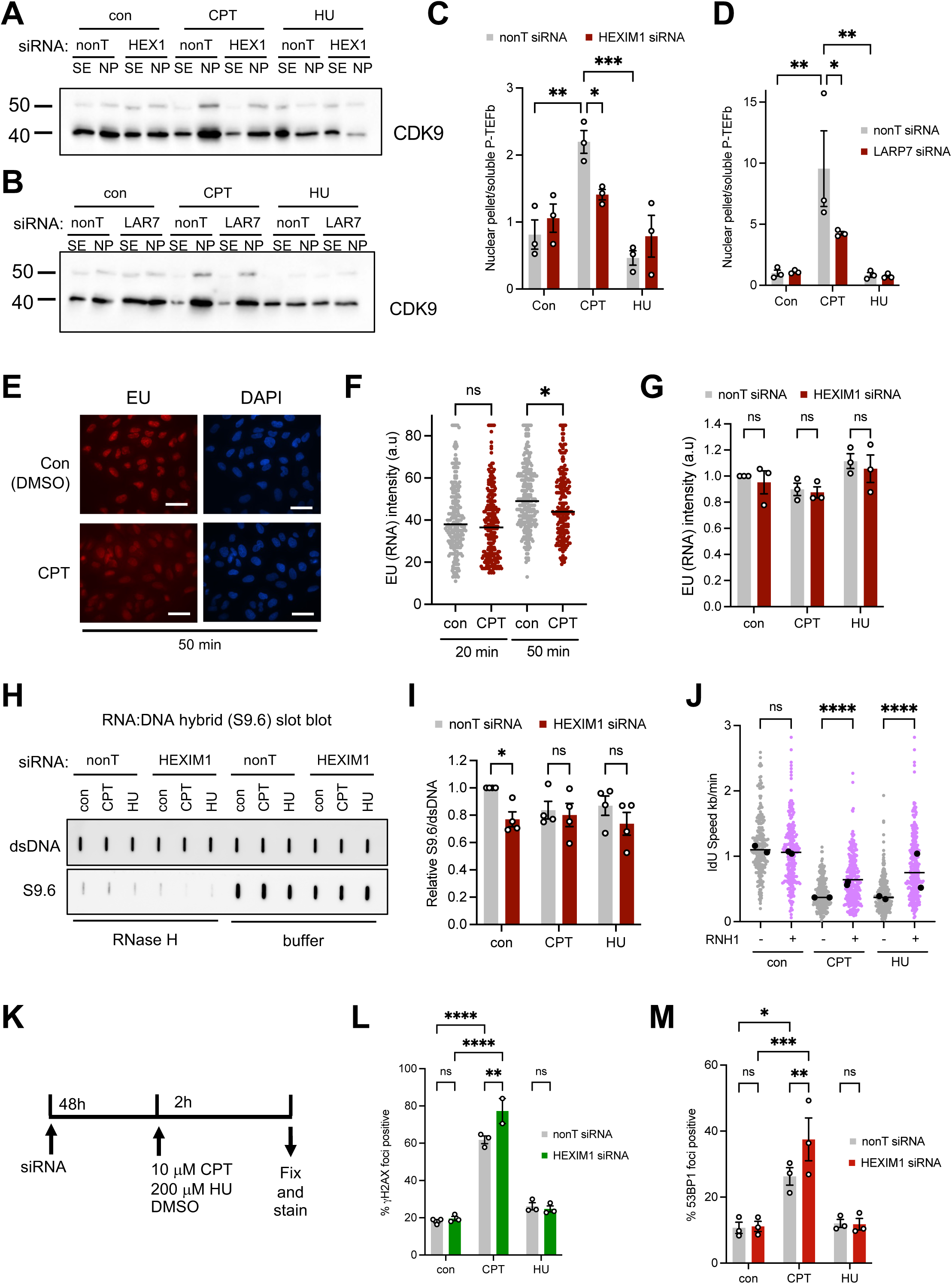
7SK-snRNP complex components do not promote transcription-replication conflicts. A) Protein levels of CDK9 in soluble extract (SE) and nuclear pellet (NP) fractions after 2 h treatment with 10 μM CPT or 200 μM HU +/- HEXIM1 siRNA (48 h). B) Protein levels of CDK9 in SE and nuclear NP fractions +/- LARP7 siRNA as in A. C) Relative levels of SE (7SK-snRNP bound) versus NP (chromatin-bound) P-TEFb after treatment +/- HEXIM1 siRNA as in A. n=3. D) Relative levels of SE versus NP P-TEFb after treatment +/- LARP7 siRNA as in B. n=3. E) Representative images of nascent RNA labelling with 5-ethynyluridine (EU). DNA was detected using DAPI. Bars: 50 μm. F) Nuclear EU intensities after treatment with 10 μM CPT and EU for 20 or 50 min. n=1. G) Nuclear EU intensities after treatment with 10 μM CPT or 200 μM HU (2 h) +/- HEXIM1 siRNA. n=3. H) Slot blots of genomic DNA stained with S9.6 antibody (RNA:DNA hybrids) and double-stranded DNA (dsDNA; loading control) after treatment with 10 μM CPT or 200 μM HU (20 min) +/- HEXIM1 siRNA. RNase H treatment was used as control. I) RNA:DNA hybrid quantification as in E. n=4. J) Replication fork speeds (IdU label) after CPT or HU treatment overexpression of turboGFP-RNase H1 (+) or eGFP vector control (-). Data from 2 repeats. K) Approach for siRNA depletion and 10 μM CPT or 200 μM HU treatment prior to immunostaining. L) Percentages of cells with >10 γH2AX foci after 2 h CPT or HU +/- HEXIM1 siRNA. n=2-3. M) Percentages of cells with >9 53BP1 foci after 2 h CPT or HU +/- HEXIM1 siRNA. n=3. Scatter blots show aggregates and medians (black points) of independent repeats with overall median (line). ANOVA or Kruskal-Wallis with multiple comparisons test, * p ≤ 0.05; ** p ≤ 0.01; *** p ≤ 0.001; **** p ≤ 0.0001; ns: not significant.

We used EU incorporation assays to test whether P-TEFb release supports increased global nascent RNA synthesis after drug treatment, which could then promote conflicts between transcription and replication. However, CPT treatment slightly decreased global levels of nascent RNA synthesis (Figure 3E, F), and HEXIM1 depletion did not further decrease global nascent RNA synthesis in cells treated with either CPT or HU (Figure 3G). This is consistent with a previous report showing limited impact of 7SK loss on global transcription after UV irradiation^7^. Next, we tested whether HEXIM1 might be required for RNA:DNA hybrid formation in response to CPT treatment. Slot blot of isolated genomic DNA using the S9.6 antibody that detects RNA:DNA hybrids ^36^ showed that HEXIM1 depletion caused a small reduction in RNA:DNA hybrid levels in untreated cells, but there was no difference when combined with CPT or HU treatment for 20 min (Figure 3H, I). We then transiently overexpressed ectopic human ribonuclease H1 (RNase H1) to reduce RNA:DNA hybrid levels, which partially rescued replication fork slowing induced both by CPT and by HU (Figure 3J, Figure S3E). This suggested that RNA:DNA hybrids are involved in replication fork slowing induced by both treatments, and not specific to CPT.

If HEXIM1 was responsible for promoting transcription-replication conflicts and thereby the source of replication stress, the HEXIM1 depletion might be expected to reduce the amount of DNA damage caused by CPT. Instead, HEXIM1 depletion slightly increased CPT-induced DNA damage as measured by γH2AX and 53BP1 nuclear foci formation (Figure 3K-M). Taken together, these data did not support that HEXIM1-induced replication fork slowing is due to HEXIM1 promoting conflicts between transcription and replication.

### HEXIM1 is required for HR repair and replication fork reversal

We had shown that CPT and HU had different impacts on transcription and activation of transcription stress responses, but these drugs also differ further in the mechanisms by which they cause replication fork slowing. Specifically, fork slowing induced by CPT, but not by HU, depends on RAD51-mediated replication fork reversal^37^. We therefore asked whether the requirement for 7SK-snRNP components in CPT-induced fork slowing might be due to a role for the 7SK-snRNP in promoting fork reversal. We first confirmed the specific requirement for fork reversal in CPT-induced fork slowing by depleting fork reversal factor ZRANB3 (Figure 4A). ZRANB3 depletion rescued replication fork slowing induced by CPT, but not by HU (Figure 4B). We next depleted fork reversal and HR factor RAD51 (Figure 4C). Because RAD51 depletion on its own slightly reduced replication fork speeds, we display IdU/CldU ratios for this experiment, and for other treatments that reduced fork speeds on their own. RAD51 depletion rescued CPT-induced fork slowing, and combination of RAD51 and HEXIM1 depletion had no additional effect on fork speeds, supporting that RAD51 and HEXIM1 may act in the same pathway (Figure 4D).

**Figure 4.**
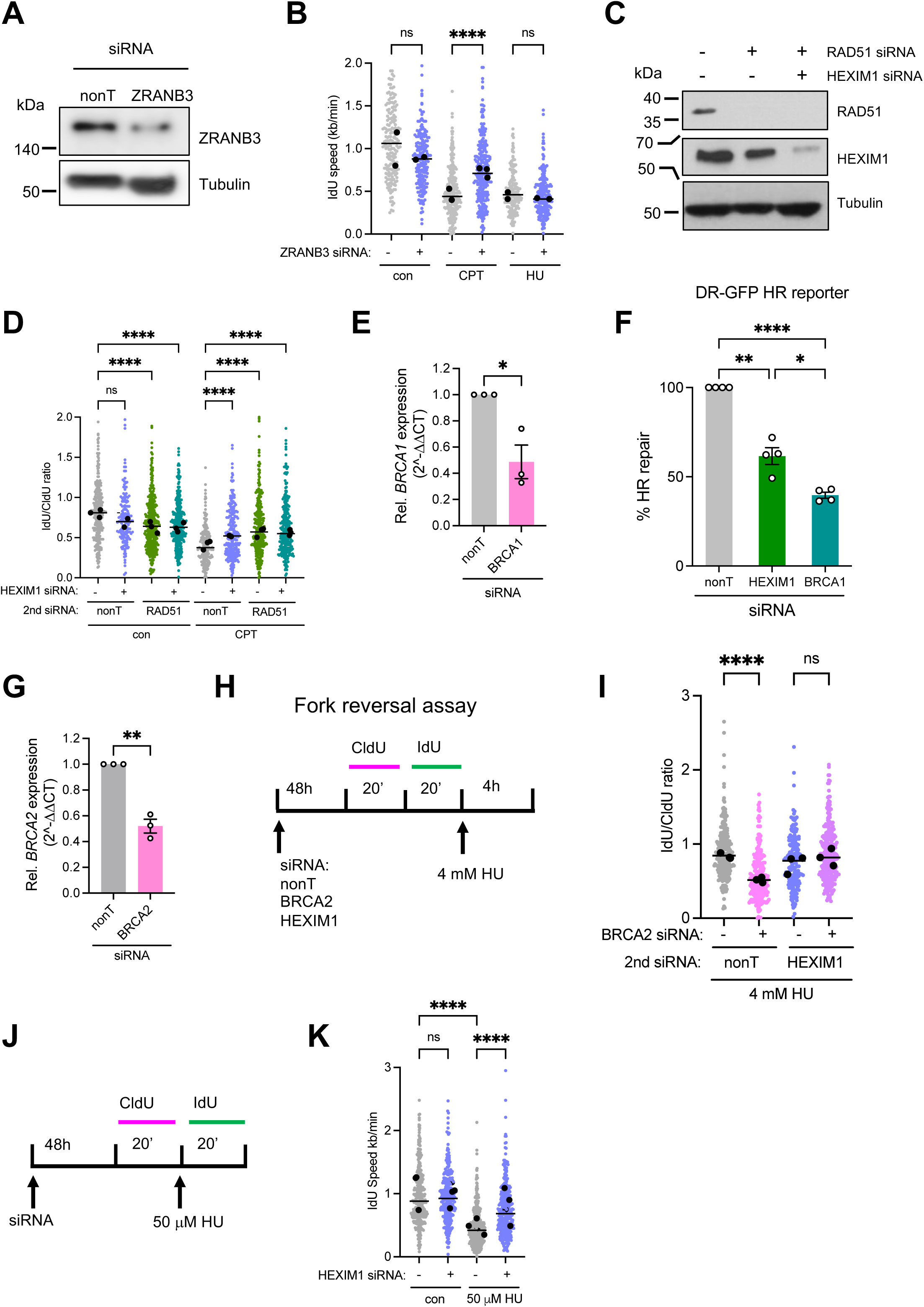
HEXIM1 promotes replication fork reversal. A) Protein levels of ZRANB3 and Tubulin (loading control) in U2OS cells 48 h after siRNA transfection. B) Replication fork speeds (IdU label) after CPT or HU treatment in presence of ZRANB3 (+) or nonT (-) siRNA. Data from 2 repeats. C) Protein levels of RAD51, HEXIM1 and Tubulin (loading control) in U2OS cells in presence of HEXIM1 (+) or nonT (-) siRNA +/- RAD51 siRNA. D) IdU/CldU ratios after CPT treatment +/- HEXIM1 and/or RAD51 siRNA. Data from 3 repeats. E) RT-qPCR of BRCA1 mRNA expression in U2OS cells 48 h after siRNA transfection. n = 3. F) Relative homologous recombination (HR) repair by gene conversion in U2OS cells harbouring a DR-GFP reporter plasmid, +/- HEXIM1 or BRCA1 siRNA. n = 4. G) RT-qPCR of BRCA2 mRNA expression in U2OS cells 48 h after siRNA transfection. n = 3. H) Approach for replication fork protection assay, using BRCA2 and/or HEXIM1 siRNA depletion prior to DNA fibre labelling followed by 4 mM HU (4 h). I) IdU/CldU ratios after 4 mM HU treatment +/- BRCA2 siRNA and/or HEXIM1 siRNA. Data from 3 repeats. J) Approach for siRNA depletion prior to DNA fibre labelling in presence of 50 μM HU. K) Replication fork speeds (IdU label) after 50 μM HU treatment in presence of HEXIM1 (+) or nonT (-) siRNA. Data from 3 repeats. Scatter blots show aggregates and medians (black points) of independent repeats with overall median (line). Kruskal-Wallis with multiple comparisons test, * p ≤ 0.05; ** p ≤ 0.01; **** p ≤ 0.0001; ns: not significant.

To test whether HEXIM1 promotes HR, we used a U2OS cell line harbouring the DR-GFP reporter construct^38^. After the rare-cutting I-SceI endonuclease induces a double-ended DSB (deDSB) in the reporter, expression of wildtype GFP shows HR repair of this site by gene conversion. HEXIM1 depletion strongly reduced HR repair in this assay, compared to BRCA1 depletion used as a positive control, without strongly changing cell cycle distribution (Figure 4E, F, Figure S4A, B). This supports that HEXIM1 promotes HR repair independently of endogenous sources of replication stress such as conflicts between transcription and replication. Similarly, MEPCE has been reported to promote RAD51 loading and HR^22^.

To investigate a role for HEXIM1 in replication fork reversal more directly, we employed a DNA fibre labelling protocol that can measure fork reversal capacity. In absence of BRCA2, reversed forks are vulnerable to nuclease degradation. This leads to shortening of the nascent labelled DNA strand during a prolonged replication block with a high dose of HU. If nascent strands are not shortened under these conditions, this is indicative of a defect in fork reversal^39^. U2OS cells were treated with siRNA against BRCA2 or HEXIM1 alone or in combination (Figure 4G, Figure S4C) and DNA fibre labelling with CldU and IdU was performed as before, but the IdU label was followed by a 4 h block with 4 mM HU (Figure 4H, I). BRCA2 depletion alone led to IdU track shortening as indicated by a decreased IdU/CldU length ratio, but when HEXIM1 was depleted, IdU tracks did not shorten either with or without BRCA2 depletion (Figure 4I). This suggests that HEXIM1 is required for replication fork reversal through a role in RAD51 function.

To investigate whether HEXIM1 is also required for fork reversal induced by low doses of HU, we treated cells with 50 μM HU, where fork slowing is mostly caused by transcription-dependent fork reversal rather than nucleotide depletion^28^. HEXIM1 depletion rescued replication fork slowing induced by 50 μM HU, suggesting that HEXIM1 is required for fork reversal even when there is no transcription stress (Figure 4J, K).

If fork reversal is prevented, then specialised DNA polymerases such as PRIMPOL or REV1 may take over and support higher replication fork speeds^40^. However, neither PRIMPOL nor REV1 appeared to be required for the increased replication fork progression in presence of CPT and absence of HEXIM1 (Figure S4D-F).

### HEXIM1 and LARP7 promote RAD51 recruitment in response to a subset of DNA damaging agents

We next used CPT, HU and JQ1 to further investigate the role of 7SK-snRNP components in HR activation in presence of different levels of transcription stress. Nuclear RAD51 foci formation was used as a measure of RAD51 recruitment to damage sites and initiation of HR (Figure 5A, B). Treatment with either 10 μM CPT or 200 μM HU for 2 h induced similar levels of RAD51 foci. Both HEXIM1 or LARP7 depletion reduced RAD51 foci induced by CPT, which causes transcription stress, but did not reduce RAD51 foci in response to HU, which had not caused transcription stress (Figure 5C, D). HEXIM1 depletion also reduced RAD51 foci formation in response to JQ1, another agent that causes transcription stress (Figure 5E). Similar results were observed in other human cancer cell lines, Hela and HCT116 (Figure S5A, B).

**Figure 5.**
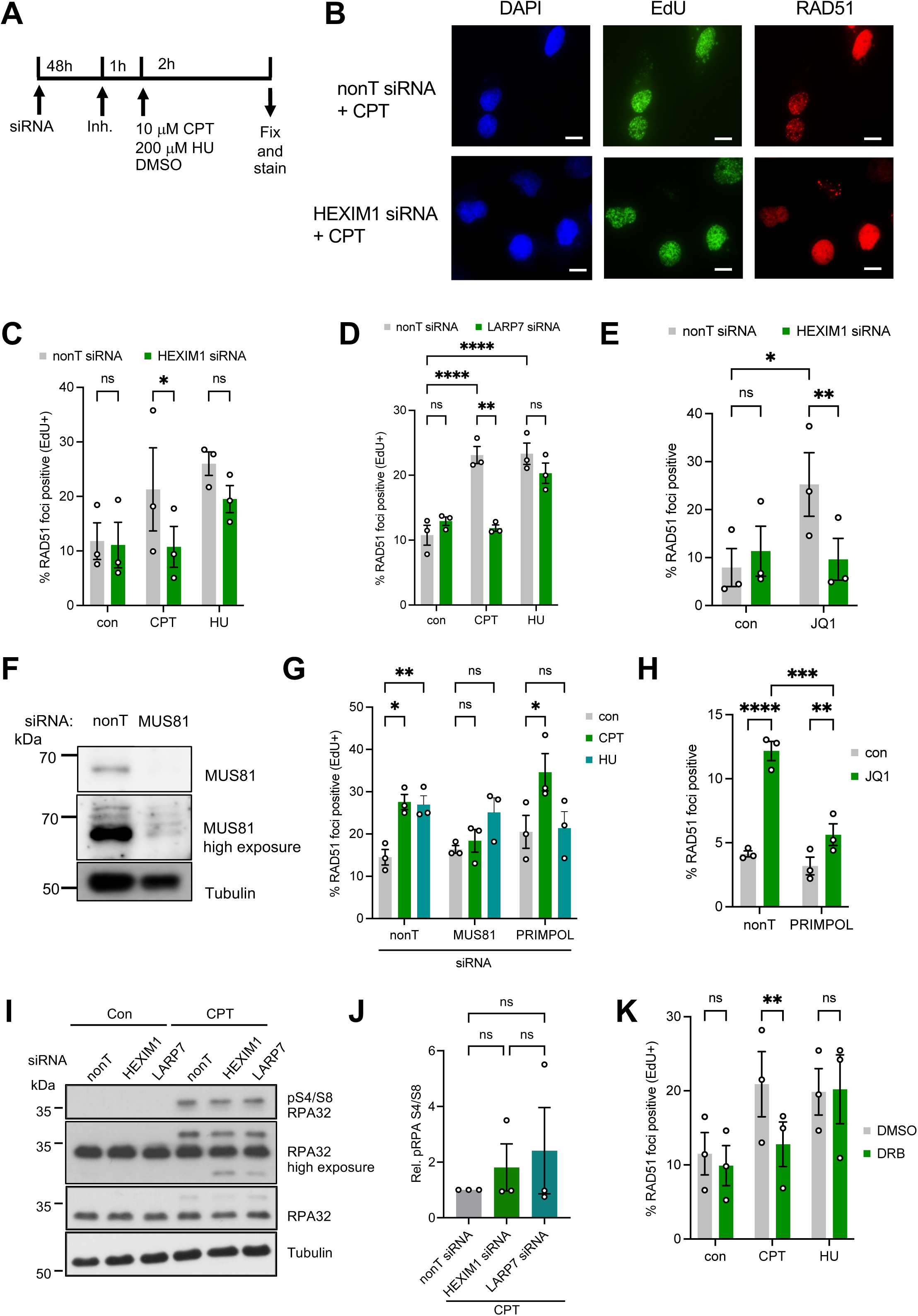
7SK-snRNP components are required for RAD51 recruitment and homologous recombination repair. A) Approach for siRNA depletion or small molecule inhibitor treatment prior to 2 h treatment with 10 μM CPT or 200 μM HU, followed by immunostaining. B) Representative images of RAD51 foci in U2OS cells after 2 h treatment with 10 μM CPT +/- HEXIM1 sRNA. C) Percentages of EdU-positive cells with RAD51 foci after CPT or HU treatment +/- HEXIM1 siRNA. n=3. D) Percentages of EdU-positive cells with RAD51 foci after CPT or HU treatment +/- LARP7 siRNA. n=3. E) Percentages of cells with RAD51 foci after 1 μM JQ1 treatment +/- HEXIM1 siRNA. n=3. F) Protein levels of MUS81 and Tubulin (loading control) 48 h after siRNA transfection. G) Percentages of EdU-positive cells with RAD51 foci after CPT or HU treatment +/- MUS81 or PRIMPOL siRNA. n=3. H) Percentages of cells with RAD51 foci after 1 μM JQ1 treatment (4 h) +/- PRIMPOL siRNA. n=3. I) Protein levels of phospho-serine 4/8 RPA32, RPA32 (loading control), and Tubulin (loading control) after 2 h 10 μM CPT treatment +/- HEXIM1 or LARP7 siRNA. J) Quantified levels of phospho-serine 4/8 RPA normalised to RPA32 after treatment as in I, relative to nonT siRNA. n=3. K) Percentages of EdU-positive cells with RAD51 foci after CPT or HU treatment +/- DRB. n=3. Column graphs show mean ± SEM. ANOVA with multiple comparisons test, * p ≤ 0.05; ** p ≤ 0.01; *** p ≤ 0.001; **** p ≤ 0.0001; ns: not significant.

We next investigated whether different lesion types underlie the differential requirements for RAD51 foci formation in response to CPT versus HU, as RAD51 can form foci either at DSBs or at post-replicative single-stranded gaps^15^. We depleted the structure-specific endonuclease MUS81 to prevent seDSB formation at replication forks^41^ (Figure 5F), and depleted PRIMPOL to prevent post-replicative gap formation. This showed that CPT-induced RAD51 foci formed at MUS81-induced seDSBs as expected, while HU- and JQ1-induced RAD51 foci both appeared to form at PRIMPOL-induced post-replicative gaps (Figure 5G, H). As HEXIM1 is required for RAD51 foci formation in response to both CPT and JQ1, this supports that 7SK-snRNP can promote RAD51 loading onto either seDSBs or post-replicative gaps. Taken together, these data suggested that while CPT, HU and JQ1 all induce DNA lesions that activate HR, the 7SK-snRP components promote RAD51 recruitment in response to those treatments that cause transcription stress.

An inability to recruit RAD51 to sites of DNA damage can result from defects in DNA end resection, and levels of phospho-Serine4/8 RPA after 1 h CPT treatment are a readout for CtIP-dependent end resection^42^. HEXIM1 or LARP7 depletion did not reduce phospho-Serine4/8 RPA formation in response to CPT (Figure 5 I-J, Figure S5C). This suggested a RAD51 recruitment defect rather than a DNA end resection defect.

DRB treatment prevented CPT-, but not HU-induced RAD51 foci formation (Figure 5K). This suggests that RAD51 recruitment in response to 200 μM HU, which does not cause transcription stress, requires neither 7K-snRNP components nor CDK9 activity or RNAPII elongation.

Because ongoing transcription favours HR repair^18,43^ and helps recruit HR factors^44^, we tested whether HEXIM1 or LARP7 depletion altered transcription of the eGFP gene after break induction. We also tested levels of antisense transcription, which could be indicative of de novo RNA synthesis at the 3’-overhang generated by resection^45,46^ (Figure S6A). In our hands, antisense transcription was not reliably increased after DSB induction, and this increase was not dependent on CtIP (resection), HEXIM1 or LARP7 (Figure S6B). While HEXIM1 or LARP7 depletion slightly altered the basal expression of eGFP mRNA, HEXIM1 or LARP7 were not required for eGFP mRNA transcription after DSB induction (Figure S6C, D). HEXIM1 or LARP7 depletion did also not prevent the rapid induction of transcription factors *EGR1* and *SERTAD1* in response to CPT (Figure S6E).

### LARP7 promotes RAD51 foci formation and replication fork slowing in a separate pathway from BRCA1

It was previously reported that LARP7 loss increased RAD51 foci formation after X-ray irradiation^20^, and two studies reported that LARP7 loss either increased or slightly decreased HR in the DR-GFP reporter construct^20,22^. This potentially suggested a separate role for LARP7 compared to other 7SK-snRNP components such as MEPCE^22^ and HEXIM1 (Figure 4F). We therefore again probed the involvement of LARP7 in HR at direct or deDSBs. In our hands, LARP7 depletion overall decreased HR in the DR-GFP reporter assay, albeit to a slightly lesser extent than HEXIM1 depletion (Figure 6A, Figure S4B). Secondly, both LARP7 and HEXIM1 depletion slightly decreased X-ray induced RAD51 foci formation (Figure 6B). Similarly, MEPCE has been reported to support RAD51 recruitment at a subset of endonuclease-induced deDSBs^22^. These data support that like HEXIM1 and MEPCE, LARP7 promotes HR, although it may be less able to promote HR repair at deDSBs.

**Figure 6.**
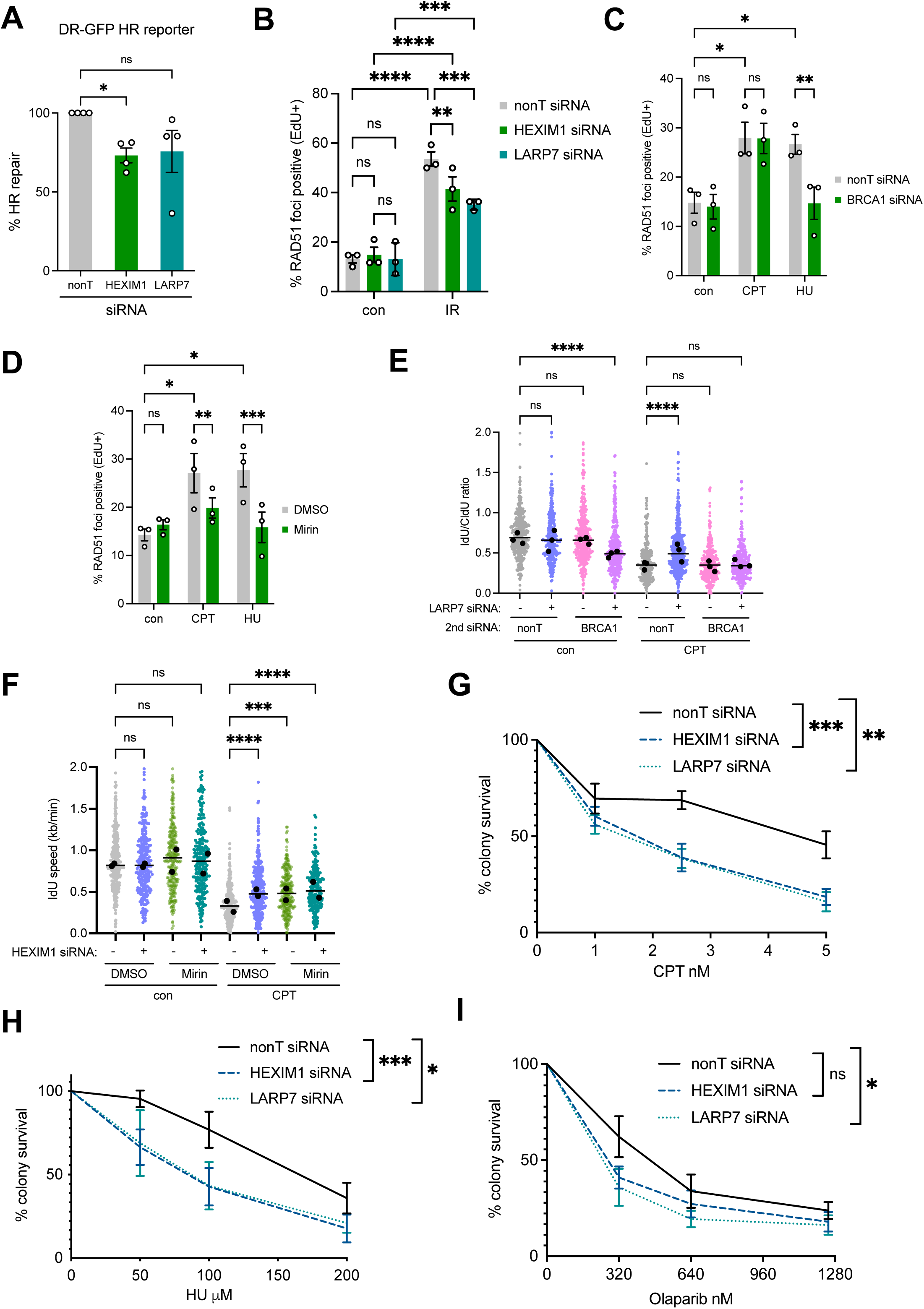
LARP7 promotes fork reversal and homologous recombination independently of BRCA1. A) Relative homologous recombination (HR) repair by gene conversion in U2OS cells harbouring a DR-GFP reporter plasmid, +/- HEXIM1 or LARP7 siRNA. n=4. B) Percentages of EdU-positive cells with RAD51 foci 4 h after 2 Gy x-ray irradiation +/- HEXIM1 or LARP7 siRNA. n=3. C) Percentages of EdU-positive cells with RAD51 foci after CPT or HU treatment +/- BRCA1 siRNA. n=3. D) Percentages of EdU-positive cells with RAD51 foci after CPT or HU treatment +/- MRE11 inhibitor (Mirin) treatment. n=3. E) IdU/CldU ratios after CPT treatment in presence of LARP7 (+) or nonT (-) siRNA +/- BRCA1 siRNA. Data from 3 repeats. F) IdU/CldU ratios after CPT treatment +/- HEXIM1 siRNA and/or MRE11 inhibitor (Mirin) treatment. n=2. G) Colony survival assay of U2OS cells after continuous treatment with CPT at the indicated concentrations +/- HEXIM or LARP7 siRNA. n=4-5. H) Colony survival assay of U2OS cells after continuous treatment with HU at the indicated concentrations +/- HEXIM or LARP7 siRNA. n=4-7. I) Colony survival assay of U2OS cells after continuous treatment with PARP inhibitor (Olaparib) at the indicated concentrations +/- HEXIM or LARP7 siRNA. n=4. Scatter blots show aggregates and medians (black points) of independent repeats with overall median (line). Column graphs show mean ± SEM. ANOVA or Kruskal-Wallis with multiple comparisons test, * p ≤ 0.05; ** p ≤ 0.01; *** p ≤ 0.001; **** p ≤ 0.0001; ns: not significant.

**Figure 7.**
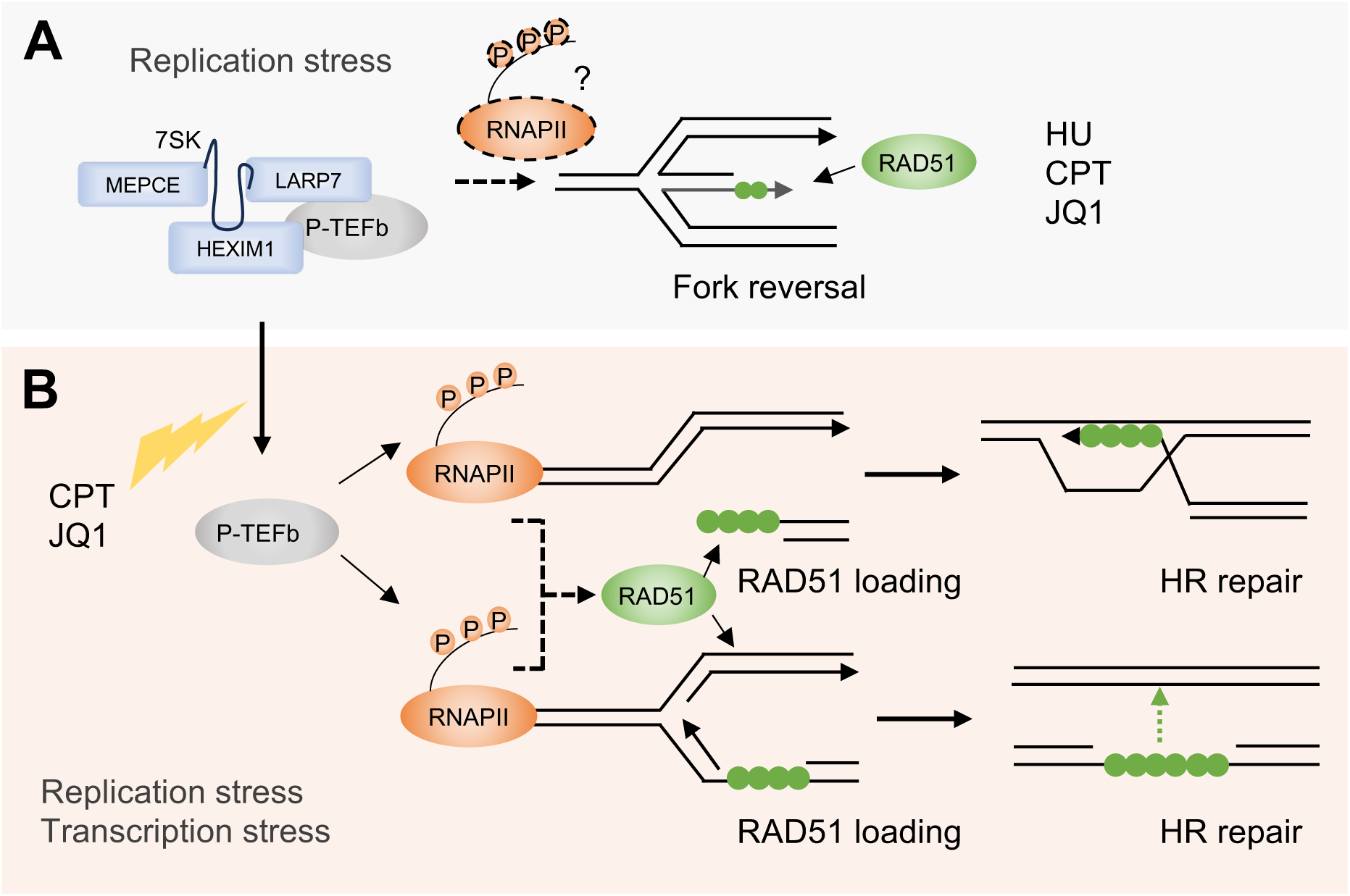
Model for the roles of the 7SK-snRNP in replication stress and homologous recombination. A) 7SK-snRNP complex components are required for replication fork reversal under all replication stress conditions, acting in the same pathway as RAD51. B) 7SK-snRNP complex components are required for RAD51 recruitment and homologous recombination repair at DSBs and post-replicative gaps specifically under conditions of transcription stress. These two scenarios could reflect separate functions of the 7SK-snRNP, or differential requirements for RNAPII activity regulation at replication forks versus DNA damage sites.

It has been suggested that LARP7 physically interacts with BRCA1, which promotes LARP7 degradation to facilitate HR^20,22^. Intriguingly, we observed that CPT-induced RAD51 foci formation appeared to not require BRCA1 (Figure 6C), although it required the activity of MRE11, which is essential for DNA end resection (Figure 6D). This agrees with a recent report that end resection at seDSBs such as those induced by CPT does not require BRCA1, with *BRCA1*^-/-^ cells displaying residual CPT-induced RAD51 foci^47^. We then combined siRNA depletion of BRCA1 and LARP7 to investigate whether LARP7 might interact with BRCA1 to rescue CPT-induced fork slowing. As reported before^21^, combined loss of BRCA1 with LARP7 caused severe replication stress in unchallenged cells (Figure 6E, Figure S7A, B). However, BRCA1 was not required for CPT-induced fork slowing, indicating that it does not act in the same pathway as LARP7 (Figure 6E). Again, CPT-induced fork slowing required MRE11 activity (Figure 6F). This raises the possibility that LARP7 can promote RAD51 functions if they are independent of BRCA1.

We had previously shown that HEXIM1 depletion sensitises U2OS cells to JQ1^19^. To probe the effect of HEXIM1 or LARP7 loss on cell survival of replication stress-inducing and DNA damaging treatments, we performed colony formation assays in presence of CPT or HU, as well as PARP inhibitor olaparib. Both HEXIM1 and LARP7 depletion did not reduce plating efficiency of untreated cells (Figure S7C), but sensitised U2OS cells to CPT (Figure 6G) and to a lesser extent to HU (Figure 6H). However, loss of HEXIM1 or LARP7 had a relatively minor impact on sensitivity to Olaparib, unlike what would be expected for canonical HR defects (Figure 6I). Taken together, this supports that 7SK-snRNP components support HR in response to a subset of treatments, and notably treatments that induce transcription stress.

## Discussion

We have shown that components of the 7SK-snRNP promote replication fork reversal, homologous recombination repair and cell survival in response to DNA damaging agents. This role becomes especially important under treatment conditions where there is evidence of transcription stress and transcriptional re-wiring.

LARP7 and MEPCE are RNA end protection factors required for 7SK RNA stability, so that LARP7 loss disrupts the entire complex^48^. In contrast, HEXIM1 is the essential regulator of the P-TEFb interaction, so that its loss would preserve the rest of the complex but abrogate its ability to control P-TEFb activity^9^. *RN7SK* knockout cells are viable, suggesting that the complex is not required for normal transcription^7^. Nevertheless, acute 7SK-snRNP loss by RNAi depletion of LARP7 can result in a baseline shift of P-TEFb toward the active state^48^, and both acute and chronic 7SK-snRNP loss prevent the DNA damage-induced release of additional P-TEFb^7^.

A sizeable fraction of HEXIM1 exists outside of the 7SK-snRNP and in a ribonucleoprotein complex with NEAT1 and DNA-PK^7,49^, and HEXIM1 promotes p53 activation^50^. LARP7 interacts with BRCA1^20^, while MEPCE was recently suggested to recruit the histone chaperone FACT complex for R-loop resolution^22^. HEXIM1 or LARP7 depletion reduced but did not abolish CDK9 recruitment to chromatin in response to high doses of CPT (Figure 3C, D). This could indicate that 7SK-snRNP is still partly functional after HEXIM1 or LARP7 depletion. However, the existence of additional yet unknown soluble P-TEFb complexes has been postulated^51^, which could contribute to CDK9 recruitment in response to stress. Nevertheless, loss of HEXIM1, LARP7 or MEPCE has similar effects on replication stress phenotypes, and all promote RAD51 loading and HR activity in several conditions. Together with the results of using DRB and p38 inhibitors, this is most consistent with a requirement for all three proteins in connection with 7SK-snRNP mediated control of P-TEFb activity rather than for their independent roles outside the complex.

The reduced CPT- and JQ1-induced fork slowing after loss of HEXIM1 or LARP7 is likely due to reduced fork reversal. Our data and data by others^22^ support that HEXIM1, LARP7 and MEPCE promote RAD51 recruitment to sites of DNA damage, which could include stressed replication forks. However, release of P-TEFb from the 7SK-snRNP appears to be only required for HR responses such as RAD51 foci formation and DSB repair, but not for fork reversal. While low-dose HU treatment does not induce signs of transcription stress, HU-induced fork reversal nevertheless depends on transcription^28^. The 7SK-snRNP could promote RAD51 recruitment to challenged replication forks via its role in regulating transcription.

In contrast, 7SK-snRNP appears to promote HR functions such as RAD51 foci formation specifically under conditions or transcription stress. In this context, transcriptional repression observed at endonuclease-induced deDSBs^52^ supports that DSBs in highly transcribed genes, such as in the DR-GFP reporter, could activate a transcription stress response. HU treatments at higher doses or for prolonged periods might also cause transcription stress: 3 mM HU stimulates CDK9 recruitment to chromatin ^53^.

The 7SK-snRNP could be required for global transcription activity, which generally favours HR repair^18^, or for HR gene expression. However, this is not consistent with the modest effects of 7SK-snRNP loss on global transcription levels after DNA damage or the fact that JQ1 treatment, while releasing P-TEFb from the 7SK-snRNP, nevertheless suppresses transcription of most protein coding genes^35^. While published data show some reduced HR gene expression in cells lacking LARP7, this was not accompanied by major changes in BRCA1, BRCA2 or RAD51 protein levels^20^.

Release of P-TEFb from the 7SK-snRNP could be required for gene transcription specifically in damaged regions. For example, ongoing transcription helps recruit HR factors to chromatin behind replication forks^44^. While transcription of protein-coding genes near endonuclease-induced deDSBs is rapidly suppressed^52^, P-TEFb could potentially counteract this repression to promote HR. After loss of 7SK RNA or MEPCE, RNAPII accumulates in gene bodies away from the transcription start site^7,21^, and this could further prevent proper regulation of RNAPII pause release in response to DNA damage. We observed that HEXIM1 or LARP7 depletion both slightly altered the expression of mRNA from the DR-GFP reporter construct (Figure S6C, D), suggesting that de-regulation of RNAPII activity could contribute to the defective HR repair of the construct.

Furthermore, recruitment of RNAPII followed by *de novo* RNA synthesis is proposed to promote HR repair at deDSBs^45,46^ and to generate RNA:DNA hybrids at stalled forks to aid fork protection^54^. *De novo* RNA synthesis at deDSBs is contested, and other studies support that RNA:DNA hybrids accumulate at deDSBs but are derived from pre-existing transcripts^43,55^. MEPCE has been implicated in resolving these RNA:DNA hybrids at a subset of deDSBs by interacting with and recruiting hybrid-resolving factors such as FACT, although increased RNA:DNA hybrids at breaks could also be a consequence of defective repair^22^. In our hands, antisense transcription was not reliably induced after deDSB induction, even though we occasionally observed increased transcript levels after depletion of HEXIM1, LARP7 or CtIP. However, any *de novo* transcripts at breaks might be transiently produced and rapidly processed, preventing reliable detection by PCR-based methods.

Finally, control of P-TEFb through 7SK-snRNP could promote specific types of RNA synthesis that regulate DNA repair factors, even away from the sites of DNA damage^56^. Finally, CDK9 itself has been implicated in promoting HR through interaction with BRCA1-BARD1^57^ and in promoting replication stress responses through an interaction with ATR, although there was no direct evidence that CDK9 promotes ATR signalling^53^.

We previously suggested that transcription-replication conflicts play a role in 7SK-snRNP-mediated fork slowing and our data cannot completely rule this out. Neither REV1 nor PRIMPOL are required for faster fork progression in absence of HEXIM1, which could support that there is not only no fork reversal but also no obstacle. However, 7SK-snRNP could help promote global fork reversal away from damage sites^58^. The differential effect of CDK9 inhibition on CPT-, versus HU-induced RAD51 foci formation also suggests that transcription is involved in generating the lesions that require RAD51 for repair. However, this need not suggest transcription-replication conflicts because CPT causes genotoxic stress by interfering with the transcription machinery directly. JQ1 treatment does not cause any known sources of replication stress except increased global RNA synthesis and R-loop accumulation^19,59,60^. Nevertheless, JQ1 treatment only induced DNA damage when HEXIM1 was depleted, supporting that even here HEXIM1’s role is to promote RAD51 recruitment and DNA repair^19^.

RNAPII activity promotes replication fork slowing, but rather than being due to transcription-replication conflicts, this could be a sign of a functional replication stress response. Future research should continue to investigate the relationship between transcription and replication stress responses. As combined loss of BRCA1 and either LARP7 or MEPCE has been reported to be synthetic lethal^21^, and HEXIM1- or LARP7-depleted cells are hypersensitive to transcription-stress inducing treatments, this could have implications for the future development of targeted cancer therapies.

## Materials and Methods

### Cell lines and reagents

U2OS cells were obtained from ATCC. HCT116 cells were obtained from Charles Swanton. U2OS-DR-GFP cells were generously provided by Jeremy Stark (City of Hope, Duarte U.S.A.). U2OS and HCT116 cells were authenticated by STR profiling (ATCC). Cells were confirmed to be free of Mycoplasma infection by PCR and grown in Dulbecco’s modified Eagle’s Medium (DMEM) with 10% foetal bovine serum (FBS), L-Glutamine (2 mM), Streptomycin (50 μg/mL) and Penicillin (50 U/mL) in a humidified atmosphere containing 5% CO_2_.

Camptothecin (CPT), hydroxyurea (HU), Hexamethylene bisacetamide (HMBA, 5 mM), 5,6-Dichlorobenzimidazole 1-β-D-ribofuranoside (DRB, 100 μM) and REV1 inhibitor JH-RE-06 (2.5 μM) were from Sigma-Aldrich. (+)-JQ1 was from Tocris Bioscience. P38 MAPK inhibitor (SB203580, Startech Scientific 10μM) was from Selleckchem. Mirin (50 μM) was from Sigma Aldrich.

### CDK9 salt fractionation

U2OS cells were harvested after treatments before being washed twice in cold PBS. Cells were lysed in low salt buffer (10 mM KCl, 10 mM MgCl_2_, 10 mM HEPES-KOH pH 7.5, 1 mM EDTA, 1 mM DTT, 0.5% Nonidet P-40, and proteinase inhibitor mixture) before being incubated on ice for 10 min. Lysates were centrifuged 5000 x g for 5 minutes and supernatant extracted which represented the soluble fraction. The pellets were washed in low salt buffer and then resuspended in high salt buffer (450 mM NaCl, 1.5 mM MgCl_2_, 20 mM HEPES pH 7.5, 0.5 mM EDTA, 1 mM DTT, 0.5% Nonidet P-40, and proteinase inhibitor mixture) and incubated on ice for 10 min. Samples were then spun down and suspensions were collected which corresponded with the nuclear pellet fraction.

### siRNA and DNA transfection

ON-TARGET siRNA SMARTpools against LARP7 (L-020996-01-0005), HEXIM1 (L-012225-01-0005), BRCA1 (L-012225-01, L-003461-00), siGenome SMARTpool against MUS81 (D-016143), siGenome siRNA against ZRANB3 (D-010025-03), custom siRNAs against RAD51^61^, CtIP^62^ and PRIMPOL^63^ were from Dharmacon/Horizon Discovery, and “Allstars negative control siRNA” was from Qiagen. 5×10^4^ cells/ml were plated in 6-well plates and transfected the next day with 50 nM siRNA using Dharmafect-1 reagent (Horizon Discovery). For RNase H1 overexpression, cells were plated in 6-well plates at 70% confluency and transfected the next day transfected using TransIT-2020 (Mirus Bio) with 2.5 μg pCMV6-AC-RNase H1-GFP (Origene) or control plasmid pEGFP-C2 (Clontech).

### Western blotting

Cells were lysed by pipetting up and down in UTB buffer (50 mM Tris-HCl pH 7.5, 150 mM β-mercaptoethanol, 8 M urea) and lysates were sonicated twice for 10 seconds each to release DNA-bound proteins. Proteins were separated by SDS-PAGE before being transferred to a nitrocellulose membrane. Primary antibodies used were monoclonal rabbit anti-HEXIM1 (Abcam ab25388,1:5000), anti-LARP7 (Abcam, ab134757,1:1000), rabbit anti-CDK9 (Cell Signalling C12F7. 1:1,000), rabbit anti-Turbo-GFP to detect GFP-RNase H1 (Evrogen AB513, 1:10,000), rabbit anti-ZRANB3 (Proteintech 23111-1-AP, 1:500), rabbit anti-RAD51 (Abcam, ab63801, 1:1,000), mouse anti-αTUBULIN (Sigma-Aldrich T6074, 1:10,000), rabbit anti-βACTIN (Cell Signaling 4967, 1:5,000). The membrane was washed before being probed with HRP-linked secondary antibodies. Detection was performed using ECL detection or ChemiDoc™ MP Imaging System (BioRad) and quantified using ImageJ.

### DNA fibre analysis

U2OS cells were pulse-labelled with 25 μM CldU and 250 μM IdU, where treatments with CPT or HU were added at the same time with IdU as indicated. Cells were harvested and DNA fibre spreads were performed mixing cell sample with spreading buffer (200 mM Tris pH 7.4, 50 mM EDTA, 0.5% SDS) to lyse cells before being spread down the slide. Fibre spreads were denatured with 2.5 M HCl for 1 h 15 min, incubated with rat anti-BrdU (BU1/75, Abcam ab6326, 1:700) and mouse anti-BrdU (B44, Becton Dickinson 347580, 1:500) for 1 h, fixed with 4% paraformaldehyde (PFA) and incubated with anti-rat AlexaFluor 594 and anti-mouse AlexaFluor 488 (Invitrogen, 1:500) for 1.5 h before mounting in Fluoroshield mounting medium (Sigma-Aldrich). Fibers were examined using a Nikon E600 microscope with a Nikon Plan Apo 60x (1.3 NA) lens, a Andor digital camera and the NIS elements acquisition software (Nikon Instruments). For quantification of fork speeds, the lengths of labelled tracks were measured using the ImageJ software^64^ (http://rsbweb.nih.gov/ij/).

### Immunofluorescence

Cells were fixed with 4% paraformaldehyde for 10 min and permeabilised with 0.25% Triton X-100 for 5 min at 4 °C followed by blocking with 2% bovine serum albumin, 0.2% Tween 20 in PBS. Primary antibodies were rabbit polyclonal anti-RAD51 (Abcam ab63801, 1:500), mouse monoclonal anti-phospho-Histone H2AX (Ser139) (JBW301, Millipore, 1:1,000), rabbit polyclonal anti-53BP1 (Bethyl A300-272A, 1:15,000). Secondary antibodies were anti-rabbit IgG AlexaFluor 594 and anti-mouse IgG AlexaFluor 488 (ThermoFisher). For RAD51 foci detection, cells were also incubated with 1 μM EdU for 30 min to mark replicating cells, followed by Click reaction according to the the Click-iT EdU Imaging Kit (ThermoFisher) manufacturer’s instructions. Coverslips were mounted in Fluoroshield with DAPI (Sigma-Aldrich). DNA was counterstained with DAPI and slides were examined using a Nikon E600 microscope as before. Cells with more than 5 RAD51 foci and 8 γH2AX or 10 53BP1 foci were scored as positive.

### EU incorporation assay

EU incorporation was performed using the Click-It RNA Alexa Flour 594 Imaging kit (Thermo Scientific). Cells were incubated with 1 mM EU as indicated, fixed and then permeabilised. Click reactions were performed according to manufacturer’s instructions, except that Hoechst staining was omitted and coverslips were instead mounted in Fluoroshield with DAPI (Sigma-Aldrich). Images were acquired using a Nikon E600 microscope and NIS elements acquisition software as before. Image J was used to generate a DAPI nuclear mask and mean AlexaFluor 594 intensity per pixel was quantified per nucleus.

### Chromatin RNA–seq

Cells were plated in equal numbers before being treated and harvested. The cells were then resuspended in cytoplasmic lysis buffer (0.15% NP-40, 10 mM Tris-HCl pH 7.0, 150 mM NaCl) before being incubated with ice for 5 min. The lysate was then poured on top of Sucrose buffer (10 mM Tris-HCl pH 7.0, 150 mM NaCl, 25% sucrose) and centrifuged at 16,000 g for 10 min. The pellet represents the nuclei and supernatant represents the cytosolic fraction which was removed and kept. The nuclei were resuspended in nuclei wash buffer (0.1% Triton X-100, 1 mM EDTA, in 1x PBS) before being centrifuged at 1500g for 1 min. The pellet was resuspended in glycerol buffer (20 mM Tris-HCl pH 8.0, 75 mM NaCl, 0.5 mM EDTA, 50% glycerol, 0.85 mM DTT), and then nuclei lysis buffer (1% NP-40, 20 mM HEPES pH 7.5, 300 mM NaCl, 1 M Urea, 0.2 mM EDTA, 1 mM DTT) was added before the sample was vortexed and put on ice. The sample was centrifuged further for 2 min at 18,500 g, and the nucleoplasm fraction was extracted by removing the supernatant. The chromatin was resuspended in chromatin resuspension buffer (25 μM α-amanitin, 50 Units SUPERaseIN, Protease inhibitors cOmplete, in 1x PBS before being stored at −80 °C. The chromatin fraction was extracted using the Zymol RNA extraction kit following manufacturer’s instructions before the RNA was quantified and stored at −80 °C.

Chromatin RNA-Seq was performed on chromatin fractions. Chromatin RNA-sequencing libraries were prepared using NEBNext Ultra II 8482 Directional RNA Library Prep kit for Illumina (New England Biolabs) and all libraries were paired-end sequencing on NovaSeq 6000 (Illumina). Sequences were first quality checked using FastQC(v0.11.9), (https://www.bioinformatics.babraham.ac.uk/projects/fastqc/) then quality filtered and trimmed using TrimGalore (v0.6.6). (https://www.bioinformatics.babraham.ac.uk/projects/trim_galore/). Genome mapping was performed with STAR (v2.7.10b)^65^ against the human reference genome (hg38 assembly). Samtools (v1.13)^65^ was used to retain only properly paired and mapped reads. Strand-specific bam files were created using samtools (https://www.htslib.org/), with strand-specific bigwig files created using BEDtools (v2.30.0)^66^ genome coverage functions. Counts for genes and genomic regions were calculated using samtools^65^ bedcoverage with GENCODE v43 gene annotation^67^. RPKM and LogFC values were calculated using R Studio (v4.4.0)^68^, with differential genes identified by LogFC > 2. Upregulated and downregulated genes were classified as those genes which were upregulated or downregulated in all 3 replicates. Gene ontology of up- and downregulated genes was performed using g:Profiler)^69^. Profile plots and heatmaps were generated using deepTools (v3.5.6)^70^.

### Reverse Transcription Quantitative real-time PCR (RT-qPCR)

Cells were harvested in TRI Reagent® (Zymo Research) and RNA extraction was performed using the Direct-zol RNA Miniprep Kit (Zymo Research), including DNase I treatment, according to the manufacturer’s instructions. RNA quality and concentration was determined using the 4200 TapeStation (Agilent) or a NanoDrop Lite spectrophotometer (ThermoFisher). Total RNA (250 ng – 1 μg) was reverse-transcribed using a mix of poly-A and random hexamer primers and the qScript cDNA synthesis kit (Quantabio) or Tetro™ cDNA synthesis kit (Meridian Bioscience), following the manufacturer’s instructions. For transcription in the DR-GFP reporter, ProtoScript® First Strand cDNA Synthesis Kit (NEB) and site-specific primers were used. Primers were from Merck (Supplementary Table S2). 2 μl cDNA and primers were added to SYBR™ Green PCR Master Mix (ThermoFisher) and analysed using the Real Time PCR QuantStudio 5 system (ThermoFisher). Cycling parameters were 95°C/5 min, 40 cycles of 95°C/15-30 seconds, annealing (temperatures see Supplementary Table S2) for 15-30 seconds and 72°C/15-30 seconds. ΔCt values were normalized to *ACTB,* or *RPLP0* for transcription in the DR-GFP reporter.

### Slot blot

Genomic DNA was extracted using the DNeasy Blood & Tissue kit (Qiagen). 10 µg genomic DNA per sample was treated with 2 U/µg DNA of RNaseH (NEB, M0297), or mock treated, for 2 hours at 37 °C. 250 ng per well was loaded onto the slot blot apparatus, transferred onto pre-wetted nylon membrane (Amersham Hybond N+). After UV crosslinking at 1200 µJ and blocking in sterile 5 % milk/TBST for 1 h at room temperature. The membrane was then incubated overnight at 4 °C in mouse anti-DNA-RNA hybrid antibody (S9.6 hybridoma growth medium, ATCC HB-8730, 1:1,000) or mouse anti-dsDNA antibody (Abcam ab27156, 1:100,000) in sterile 5 % BSA/TBST, before being washed in TBST and incubated in goat anti-mouse HRP (Cell Signaling 7074, 1:5,000) 5 % milk/TBST for 1 hour at room temperature. Images were collected using a ChemiDoc™ MP Imaging System (BioRad) and quantified using ImageJ.

### Homologous recombination assay

U2OS-DR-GFP cells and expression constructs for RFP and I-Sce I endonuclease were generously provided by Jeremy Stark (City of Hope, Duarte U.S.A.). U2OS-DR3-GFP cells were transfected with the indicated siRNAs using Dharmafect1 (Dharmacon). After 24 h, growth medium was replaced, and cells were co-transfected with RFP and I-Sce I constructs using FuGene6 (Promega) or *Trans*IT^®^-2020 (Mirus). 24 h later, the medium was replaced and cells were cultured for an additional 48 h before fixation in 2% PFA in PBS. GFP-positive cells were detected on a Fortessa X-20 flow cytometer and analysed with the BD FacsDiva software. A total of 20,000 cells were acquired per sample, and the percentage of GFP-positive cells was determined as a fraction of RFP positive cells.

### Propidium iodide flow cytometry

Cells were fixed in 70% ethanol at −20 °C for 30 min, then centrifuged at 500 g for 5 min. After two washes in PBS, cells were mixed with PBS containing 10 µg/ml propidium iodide and 25 µg/ml ribonuclease A (Merck). Cell cycle profiles were generated and analysed using the Fortessa X-20 flow cytometer and BD FacsDiva software, respectively.

### Colony survival assay

48 h after siRNA transfection, U2OS cells were trypsinised and plated in triplicate at a density of 800-2000 cells per well. Cells were treated with drugs continuously until colonies of >50 cells had grown, then stained with 50% ethanol, 2% methylene blue and colonies counted. IC50 values were calculated using the built-in nonlinear regression (curve fit) function in GraphPad Prism. The equation used was [Inhibitor] vs. normalized response -- Variable slope, least squares fit.

### Quantification and statistical analysis

Values represent the means + 1x SEM of at least 3 independent biological repeats. The number of independent biological repeats (n) is indicated in the figure legends. For foci analysis, at least 10 different areas were quantified for each independent biological repeat. Gaussian distribution was determined and statistical tests were performed using the GraphPad Prism 10 software, version 10.2.0 (392). For comparing two datasets with non-Gaussian distribution, Mann-Whitney test was used. For multiple comparisons, one-or two-way ANOVA or mixed-effects analysis with Dunnett’s, Tukey’s or Sidak’s test was used for datasets with Gaussian distribution and Kruskal-Wallis test with Dunn’s test for datasets with non-Gaussian distribution.

### Data and software availability

RNA sequencing data generated and analysed during this study are available in the GEO repository under accession number GSE309660, https://www.ncbi.nlm.nih.gov/geo/query/acc.cgi?acc=GSE309660.

## Acknowledgements

A.B., C.W., R.D.W.K. and R.J.W. were supported by a Cancer Research UK Programme Foundation Award to E.P. and A.K. (C25526/A28275). C.S.V. was supported by a Medical Research Council grant to E.P (MR/W031442/1). A.K.W. was supported by an EPSRC grant to J.R.M (EP/X023087/1). The authors would like to acknowledge the Genomics Birmingham Facility and Matthew Newbould at the University of Birmingham for the generation of the chromatin RNA-seq data.

## Author contributions

A.B. and E.P. conceived the study; C.W designed bioinformatics pipelines and analysed experimental data; A.B., R.D.W.K, A.K.W. and E.P. designed experiments; A.B., R.D.W.K, R.J.W., C.S.V., A.K.W. and S.M.H. performed experiments and analysed experimental data; J.R.M., A.K. and E.P. supervised the study; E.P. wrote the paper.

## References

1. Groelly, F.J., Fawkes, M., Dagg, R.A., Blackford, A.N., and Tarsounas, M. (2023). Targeting DNA damage response pathways in cancer. Nat Rev Cancer 23, 78–94. 10.1038/s41568-022-00535-5.

2. Noe Gonzalez, M., Blears, D., and Svejstrup, J.Q. (2021). Causes and consequences of RNA polymerase II stalling during transcript elongation. Nat Rev Mol Cell Biol 22, 3–21. 10.1038/s41580-020-00308-8.

3. Cybulla, E., and Vindigni, A. (2023). Leveraging the replication stress response to optimize cancer therapy. Nat Rev Cancer 23, 6–24. 10.1038/s41568-022-00518-6.

4. Fujinaga, K., Luo, Z., Schaufele, F., and Peterlin, B.M. (2015). Visualization of positive transcription elongation factor b (P-TEFb) activation in living cells. J Biol Chem 290, 1829–1836. 10.1074/jbc.M114.605816.

5. Amente, S., Gargano, B., Napolitano, G., Lania, L., and Majello, B. (2009). Camptothecin releases P-TEFb from the inactive 7SK snRNP complex. Cell Cycle 8, 1249–1255. 10.4161/cc.8.8.8286.

6. Fang, Y., Wang, Y., Spector, B.M., Xiao, X., Yang, C., Li, P., Yuan, Y., Ding, P., Xiao, Z.X., Zhang, P., et al. (2022). Dynamic regulation of P-TEFb by 7SK snRNP is integral to the DNA damage response to regulate chemotherapy sensitivity. iScience 25, 104844. 10.1016/j.isci.2022.104844.

7. Studniarek, C., Tellier, M., Martin, P.G.P., Murphy, S., Kiss, T., and Egloff, S. (2021). The 7SK/P-TEFb snRNP controls ultraviolet radiation-induced transcriptional reprogramming. Cell Rep 35, 108965. 10.1016/j.celrep.2021.108965.

8. Bugai, A., Quaresma, A.J.C., Friedel, C.C., Lenasi, T., Duster, R., Sibley, C.R., Fujinaga, K., Kukanja, P., Hennig, T., Blasius, M., et al. (2019). P-TEFb Activation by RBM7 Shapes a Pro-survival Transcriptional Response to Genotoxic Stress. Mol Cell 74, 254–267 e210. 10.1016/j.molcel.2019.01.033.

9. Quaresma, C.A.J., Bugai, A., and Barboric, M. (2016). Cracking the control of RNA polymerase II elongation by 7SK snRNP and P-TEFb. Nucleic Acids Res 44, 7527–7539 10.1093/nar/gkw585.

10. Nguyen, V.T., Kiss, T., Michels, A.A., and Bensaude, O. (2001). 7SK small nuclear RNA binds to and inhibits the activity of CDK9/cyclin T complexes. Nature 414, 322. 10.1038/35104581.

11. Fujinaga, K., Huang, F., and Peterlin, B.M. (2023). P-TEFb: The master regulator of transcription elongation. Mol Cell 83, 393–403. 10.1016/j.molcel.2022.12.006.

12. Ji, X., Lu, H., Zhou, Q., and Luo, K. (2014). LARP7 suppresses P-TEFb activity to inhibit breast cancer progression and metastasis. eLife 3, e02907. 10.7554/eLife.02907.

13. Puidebat, O., and Egloff, S. (2025). The 7SK snRNP complex: a critical regulator in carcinogenesis. Biochimie. 10.1016/j.biochi.2025.05.003.

14. Petermann, E., Orta, M.L., Issaeva, N., Schultz, N., and Helleday, T. (2010). Hydroxyurea-stalled replication forks become progressively inactivated and require two different RAD51-mediated pathways for restart and repair. Mol Cell 37, 492–502. 10.1016/j.molcel.2010.01.021.

15. Piberger, A.L., Bowry, A., Kelly, R.D.W., Walker, A.K., Gonzalez-Acosta, D., Bailey, L.J., Doherty, A.J., Mendez, J., Morris, J.R., Bryant, H.E., and Petermann, E. (2020). PrimPol-dependent single-stranded gap formation mediates homologous recombination at bulky DNA adducts. Nat Commun 11, 5863. 10.1038/s41467-020-19570-7.

16. Bhowmick, R., Lerdrup, M., Gadi, S.A., Rossetti, G.G., Singh, M.I., Liu, Y., Halazonetis, T.D., and Hickson, I.D. (2022). RAD51 protects human cells from transcription-replication conflicts. Mol Cell 82, 3366–3381 e3369. 10.1016/j.molcel.2022.07.010.

17. Groelly, F.J., Dagg, R.A., Petropoulos, M., Rossetti, G.G., Prasad, B., Panagopoulos, A., Paulsen, T., Karamichali, A., Jones, S.E., Ochs, F., et al. (2022). Mitotic DNA synthesis is caused by transcription-replication conflicts in BRCA2-deficient cells. Mol Cell 82, 3382–3397 e3387. 10.1016/j.molcel.2022.07.011.

18. Aymard, F., Bugler, B., Schmidt, C.K., Guillou, E., Caron, P., Briois, S., Iacovoni, J.S., Daburon, V., Miller, K.M., Jackson, S.P., and Legube, G. (2014). Transcriptionally active chromatin recruits homologous recombination at DNA double-strand breaks. Nat Struct Mol Biol 21, 366–374. 10.1038/nsmb.2796.

19. Bowry, A., Piberger, A.L., Rojas, P., Saponaro, M., and Petermann, E. (2018). BET inhibition induces HEXIM1- and RAD51-dependent conflicts between transcription and replication. Cell Rep 25, 2061–2069.

20. Zhang, F., Yan, P., Yu, H., Le, H., Li, Z., Chen, J., Liang, X., Wang, S., Wei, W., Liu, L., et al. (2020). LARP7 Is a BRCA1 Ubiquitinase Substrate and Regulates Genome Stability and Tumorigenesis. Cell Rep 32, 108058. 10.1016/j.celrep.2020.108058.

21. Patel, P.S., Algouneh, A., Krishnan, R., Reynolds, J.J., Nixon, K.C.J., Hao, J., Lee, J., Feng, Y., Fozil, C., Stanic, M., et al. (2023). Excessive transcription-replication conflicts are a vulnerability of BRCA1-mutant cancers. Nucleic Acids Res 51, 4341–4362. 10.1093/nar/gkad172.

22. Devanathan, S.K., Li, Y.R., Shelton, S.B., Nguyen, J., Tseng, W.C., Shah, N.M., Mercado, M., Miller, K.M., and Xhemalce, B. (2025). MePCE promotes homologous recombination through coordinating R-loop resolution at DNA double-stranded breaks. Cell Rep 44, 115740. 10.1016/j.celrep.2025.115740.

23. Yamauchi, T., Yoshida, A., and Ueda, T. (2011). Camptothecin induces DNA strand breaks and is cytotoxic in stimulated normal lymphocytes. Oncol Rep 25, 347–352. 10.3892/or.2010.1100.

24. Kouzine, F., Gupta, A., Baranello, L., Wojtowicz, D., Ben-Aissa, K., Liu, J., Przytycka, T.M., and Levens, D. (2013). Transcription-dependent dynamic supercoiling is a short-range genomic force. Nat Struct Mol Biol 20, 396–403. 10.1038/nsmb.2517.

25. Baranello, L., Wojtowicz, D., Cui, K., Devaiah, B.N., Chung, H.J., Chan-Salis, K.Y., Guha, R., Wilson, K., Zhang, X., Zhang, H., et al. (2016). RNA Polymerase II Regulates Topoisomerase 1 Activity to Favor Efficient Transcription. Cell 165, 357–371. 10.1016/j.cell.2016.02.036.

26. Duardo, R.C., Marinello, J., Russo, M., Morelli, S., Pepe, S., Guerra, F., Gomez-Gonzalez, B., Aguilera, A., and Capranico, G. (2024). Human DNA topoisomerase I poisoning causes R loop-mediated genome instability attenuated by transcription factor IIS. Sci Adv 10, eadm8196. 10.1126/sciadv.adm8196.

27. Snyder, R.D. (1984). Deoxyribonucleoside triphosphate pools in human diploid fibroblasts and their modulation by hydroxyurea and deoxynucleosides. Biochem Pharmacol 33, 1515–1518.

28. Andrs, M., Stoy, H., Boleslavska, B., Chappidi, N., Kanagaraj, R., Nascakova, Z., Menon, S., Rao, S., Oravetzova, A., Dobrovolna, J., et al. (2023). Excessive reactive oxygen species induce transcription-dependent replication stress. Nat Commun 14, 1791. 10.1038/s41467-023-37341-y.

29. Contreras, X., Barboric, M., Lenasi, T., and Peterlin, B.M. (2007). HMBA Releases P-TEFb from HEXIM1 and 7SK snRNA via PI3K/Akt and Activates HIV Transcription. PLOS Pathogens 3, e146. 10.1371/journal.ppat.0030146.

30. Sirbu, B.M., McDonald, W.H., Dungrawala, H., Badu-Nkansah, A., Kavanaugh, G.M., Chen, Y., Tabb, D.L., and Cortez, D. (2013). Identification of proteins at active, stalled, and collapsed replication forks using isolation of proteins on nascent DNA (iPOND) coupled with mass spectrometry. J Biol Chem 288, 31458–31467. 10.1074/jbc.M113.511337.

31. Nakamura, K., Kustatscher, G., Alabert, C., Hodl, M., Forne, I., Volker-Albert, M., Satpathy, S., Beyer, T.E., Mailand, N., Choudhary, C., et al. (2021). Proteome dynamics at broken replication forks reveal a distinct ATM-directed repair response suppressing DNA double-strand break ubiquitination. Mol Cell 81, 1084–1099 e1086. 10.1016/j.molcel.2020.12.025.

32. Biglione, S., Byers, S.A., Price, J.P., Nguyen, V.T., Bensaude, O., Price, D.H., and Maury, W. (2007). Inhibition of HIV-1 replication by P-TEFb inhibitors DRB, seliciclib and flavopiridol correlates with release of free P-TEFb from the large, inactive form of the complex. Retrovirology 4, 47. 10.1186/1742-4690-4-47.

33. Mayer, A., di Iulio, J., Maleri, S., Eser, U., Vierstra, J., Reynolds, A., Sandstrom, R., Stamatoyannopoulos, J.A., and Churchman, L.S. (2015). Native elongating transcript sequencing reveals human transcriptional activity at nucleotide resolution. Cell 161, 541–554. 10.1016/j.cell.2015.03.010.

34. Veloso, A., Biewen, B., Paulsen, M.T., Berg, N., Carmo de Andrade Lima, L., Prasad, J., Bedi, K., Magnuson, B., Wilson, T.E., and Ljungman, M. (2013). Genome-wide transcriptional effects of the anti-cancer agent camptothecin. PLoS One 8, e78190. 10.1371/journal.pone.0078190.

35. Muhar, M., Ebert, A., Neumann, T., Umkehrer, C., Jude, J., Wieshofer, C., Rescheneder, P., Lipp, J.J., Herzog, V.A., Reichholf, B., et al. (2018). SLAM-seq defines direct gene-regulatory functions of the BRD4-MYC axis. Science 360, 800–805. 10.1126/science.aao2793.

36. Boguslawski, S.J., Smith, D.E., Michalak, M.A., Mickelson, K.E., Yehle, C.O., Patterson, W.L., and Carrico, R.J. (1986). Characterization of monoclonal antibody to DNA.RNA and its application to immunodetection of hybrids. J Immunol Methods 89, 123–130.

37. Zellweger, R., Dalcher, D., Mutreja, K., Berti, M., Schmid, J.A., Herrador, R., Vindigni, A., and Lopes, M. (2015). Rad51-mediated replication fork reversal is a global response to genotoxic treatments in human cells. J Cell Biol 208, 563–579. 10.1083/jcb.201406099.

38. Pierce, A.J., Johnson, R.D., Thompson, L.H., and Jasin, M. (1999). XRCC3 promotes homology-directed repair of DNA damage in mammalian cells. Genes Dev 13, 2633–2638.

39. Mijic, S., Zellweger, R., Chappidi, N., Berti, M., Jacobs, K., Mutreja, K., Ursich, S., Ray Chaudhuri, A., Nussenzweig, A., Janscak, P., and Lopes, M. (2017). Replication fork reversal triggers fork degradation in BRCA2-defective cells. Nat Commun 8, 859. 10.1038/s41467-017-01164-5.

40. Bai, G., Kermi, C., Stoy, H., Schiltz, C.J., Bacal, J., Zaino, A.M., Hadden, M.K., Eichman, B.F., Lopes, M., and Cimprich, K.A. (2020). HLTF Promotes Fork Reversal, Limiting Replication Stress Resistance and Preventing Multiple Mechanisms of Unrestrained DNA Synthesis. Mol Cell 78, 1237–1251 e1237. 10.1016/j.molcel.2020.04.031.

41. Regairaz, M., Zhang, Y.W., Fu, H., Agama, K.K., Tata, N., Agrawal, S., Aladjem, M.I., and Pommier, Y. (2011). Mus81-mediated DNA cleavage resolves replication forks stalled by topoisomerase I-DNA complexes. J Cell Biol 195, 739–749. 10.1083/jcb.201104003.

42. Sartori, A.A., Lukas, C., Coates, J., Mistrik, M., Fu, S., Bartek, J., Baer, R., Lukas, J., and Jackson, S.P. (2007). Human CtIP promotes DNA end resection. Nature 450, 509–514. 10.1038/nature06337.

43. Ouyang, J., Yadav, T., Zhang, J.M., Yang, H., Rheinbay, E., Guo, H., Haber, D.A., Lan, L., and Zou, L. (2021). RNA transcripts stimulate homologous recombination by forming DR-loops. Nature 594, 283–288. 10.1038/s41586-021-03538-8.

44. Bandau, S., Alvarez, V., Jiang, H., Graff, S., Sundaramoorthy, R., Gierlinski, M., Toman, M., Owen-Hughes, T., Sidoli, S., Lamond, A., and Alabert, C. (2024). RNA polymerase II promotes the organization of chromatin following DNA replication. EMBO Rep 25, 1387–1414. 10.1038/s44319-024-00085-x.

45. D’Alessandro, G., Whelan, D.R., Howard, S.M., Vitelli, V., Renaudin, X., Adamowicz, M., Iannelli, F., Jones-Weinert, C.W., Lee, M., Matti, V., et al. (2018). BRCA2 controls DNA:RNA hybrid level at DSBs by mediating RNase H2 recruitment. Nat Commun 9, 5376. 10.1038/s41467-018-07799-2.

46. Liu, S., Hua, Y., Wang, J., Li, L., Yuan, J., Zhang, B., Wang, Z., Ji, J., and Kong, D. (2021). RNA polymerase III is required for the repair of DNA double-strand breaks by homologous recombination. Cell 184, 1314–1329 e1310. 10.1016/j.cell.2021.01.048.

47. Pavani, R., Tripathi, V., Vrtis, K.B., Zong, D., Chari, R., Callen, E., Pankajam, A.V., Zhen, G., Matos-Rodrigues, G., Yang, J., et al. (2024). Structure and repair of replication-coupled DNA breaks. Science 385, eado3867. 10.1126/science.ado3867.

48. He, N., Jahchan, N.S., Hong, E., Li, Q., Bayfield, M.A., Maraia, R.J., Luo, K., and Zhou, Q. (2008). A La-related protein modulates 7SK snRNP integrity to suppress P-TEFb-dependent transcriptional elongation and tumorigenesis. Mol Cell 29, 588–599. 10.1016/j.molcel.2008.01.003.

49. Morchikh, M., Cribier, A., Raffel, R., Amraoui, S., Cau, J., Severac, D., Dubois, E., Schwartz, O., Bennasser, Y., and Benkirane, M. (2017). HEXIM1 and NEAT1 Long Non-coding RNA Form a Multi-subunit Complex that Regulates DNA-Mediated Innate Immune Response. Mol Cell 67, 387–399 e385. 10.1016/j.molcel.2017.06.020.

50. Lew, Q.J., Chia, Y.L., Chu, K.L., Lam, Y.T., Gurumurthy, M., Xu, S., Lam, K.P., Cheong, N., and Chao, S.H. (2012). Identification of HEXIM1 as a positive regulator of p53. J Biol Chem 287, 36443–36454. 10.1074/jbc.M112.374157.

51. Graham, T.G.W., Dugast-Darzacq, C., Dailey, G.M., Weng, B., Anantakrishnan, S., Darzacq, X., and Tjian, R. (2025). Single-molecule live imaging of subunit interactions and exchange within cellular regulatory complexes. Mol Cell 85, 2854–2868 e2857. 10.1016/j.molcel.2025.06.028.

52. Awwad, S.W., Abu-Zhayia, E.R., Guttmann-Raviv, N., and Ayoub, N. (2017). NELF-E is recruited to DNA double-strand break sites to promote transcriptional repression and repair. EMBO Rep 18, 745–764. 10.15252/embr.201643191.

53. Yu, D.S., Zhao, R., Hsu, E.L., Cayer, J., Ye, F., Guo, Y., Shyr, Y., and Cortez, D. (2010). Cyclin-dependent kinase 9-cyclin K functions in the replication stress response. EMBO Rep 11, 876–882. 10.1038/embor.2010.153.

54. Song, L., Xie, H., Fan, H., Zhang, Y., Cheng, Z., Chen, J., Guo, Y., Zhang, S., Zhou, X., Li, Z., et al. (2025). Dynamic control of RNA-DNA hybrid formation orchestrates DNA2 activation at stalled forks by RNAPII and DDX39A. Mol Cell 85, 506–522 e507. 10.1016/j.molcel.2024.11.034.

55. Saur, F., Lesage, E., Pradel, L., Collins, S., Finoux, A.L., Alghoul, E., Le Bozec, B., Rocher, V., Carette, R., Puget, N., et al. (2025). Transcriptional repression facilitates RNA:DNA hybrid accumulation at DNA double-strand breaks. Nat Cell Biol 27, 992–1005. 10.1038/s41556-025-01669-y.

56. Audoynaud, C., Vagner, S., and Lambert, S. (2021). Non-homologous end-joining at challenged replication forks: an RNA connection? Trends Genet 37, 973–985. 10.1016/j.tig.2021.06.010.

57. Nepomuceno, T.C., Fernandes, V.C., Gomes, T.T., Carvalho, R.S., Suarez-Kurtz, G., Monteiro, A.N., and Carvalho, M.A. (2017). BRCA1 recruitment to damaged DNA sites is dependent on CDK9. Cell Cycle 16, 665–672. 10.1080/15384101.2017.1295177.

58. Mutreja, K., Krietsch, J., Hess, J., Ursich, S., Berti, M., Roessler, F.K., Zellweger, R., Patra, M., Gasser, G., and Lopes, M. (2018). ATR-Mediated Global Fork Slowing and Reversal Assist Fork Traverse and Prevent Chromosomal Breakage at DNA Interstrand Cross-Links. Cell Rep 24, 2629–2642 e2625. 10.1016/j.celrep.2018.08.019.

59. Kim, J.J., Lee, S.Y., Gong, F., Battenhouse, A.M., Boutz, D.R., Bashyal, A., Refvik, S.T., Chiang, C.M., Xhemalce, B., Paull, T.T., et al. (2019). Systematic bromodomain protein screens identify homologous recombination and R-loop suppression pathways involved in genome integrity. Genes Dev 33, 1751–1774. 10.1101/gad.331231.119.

60. Lam, F.C., Kong, Y.W., Huang, Q., Vu Han, T.-L., Maffa, A.D., Kasper, E.M., and Yaffe, M.B. (2020). BRD4 prevents the accumulation of R-loops and protects against transcription–replication collision events and DNA damage. Nature communications 11, 4083. 10.1038/s41467-020-17503-y.

61. Ito, M., Yamamoto, S., Nimura, K., Hiraoka, K., Tamai, K., and Kaneda, Y. (2005). Rad51 siRNA delivered by HVJ envelope vector enhances the anti-cancer effect of cisplatin. The journal of gene medicine 7, 1044–1052. 10.1002/jgm.753.

62. Yu, X., and Chen, J. (2004). DNA damage-induced cell cycle checkpoint control requires CtIP, a phosphorylation-dependent binding partner of BRCA1 C-terminal domains. Mol Cell Biol 24, 9478–9486. 10.1128/MCB.24.21.9478-9486.2004.

63. Bianchi, J., Rudd, S.G., Jozwiakowski, S.K., Bailey, L.J., Soura, V., Taylor, E., Stevanovic, I., Green, A.J., Stracker, T.H., Lindsay, H.D., and Doherty, A.J. (2013). PrimPol bypasses UV photoproducts during eukaryotic chromosomal DNA replication. Mol Cell 52, 566–573. 10.1016/j.molcel.2013.10.035.

64. Schneider, C.A., Rasband, W.S., and Eliceiri, K.W. (2012). NIH Image to ImageJ: 25 years of image analysis. Nat Methods 9, 671–675.

65. Dobin, A., Davis, C.A., Schlesinger, F., Drenkow, J., Zaleski, C., Jha, S., Batut, P., Chaisson, M., and Gingeras, T.R. (2013). STAR: ultrafast universal RNA-seq aligner. Bioinformatics 29, 15–21. 10.1093/bioinformatics/bts635.

66. Quinlan, A.R., and Hall, I.M. (2010). BEDTools: a flexible suite of utilities for comparing genomic features. Bioinformatics 26, 841–842. 10.1093/bioinformatics/btq033.

67. Mudge, J.M., Carbonell-Sala, S., Diekhans, M., Martinez, J.G., Hunt, T., Jungreis, I., Loveland, J.E., Arnan, C., Barnes, I., Bennett, R., et al. (2025). GENCODE 2025: reference gene annotation for human and mouse. Nucleic Acids Res 53, D966–D975. 10.1093/nar/gkae1078.

68. Team, R.C. (2021). R: A language and environment for statistical computing. R Foundation for Statistical Computing, Vienna, Austria.

69. Raudvere, U., Kolberg, L., Kuzmin, I., Arak, T., Adler, P., Peterson, H., and Vilo, J. (2019). g:Profiler: a web server for functional enrichment analysis and conversions of gene lists (2019 update). Nucleic Acids Res 47, W191–W198. 10.1093/nar/gkz369.

70. Ramirez, F., Ryan, D.P., Gruning, B., Bhardwaj, V., Kilpert, F., Richter, A.S., Heyne, S., Dundar, F., and Manke, T. (2016). deepTools2: a next generation web server for deep-sequencing data analysis. Nucleic Acids Res 44, W160–165. 10.1093/nar/gkw257.

