## Supplemental Data for "The 7SK small nuclear ribonucleoprotein links the cell responses to transcription and replication stress by promoting replication fork reversal and homologous recombination"

### Supplemental Figures

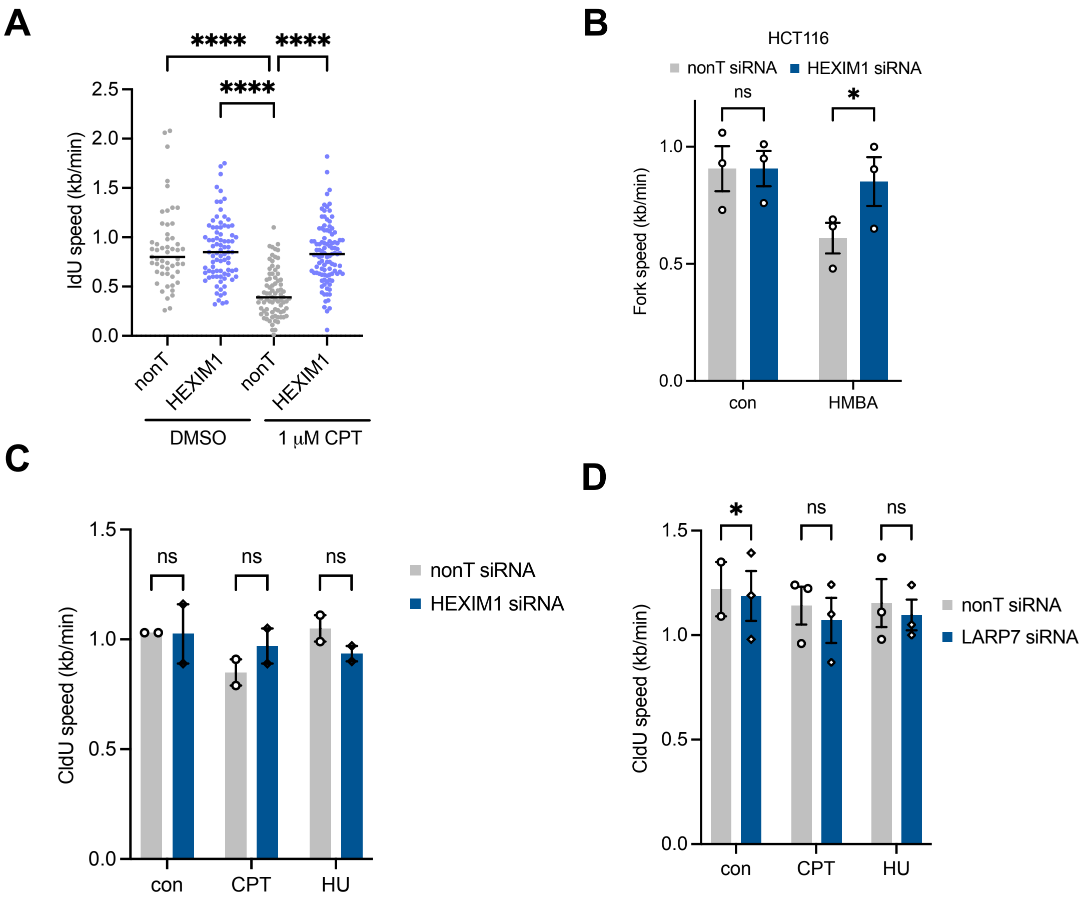

**Figure S1. 7SK-snRP component depletion rescues replication fork slowing.**
A) Second label (IdU) speeds in U2OS cells treated with non-targeting (nonT) or HEXIM1 siRNA for 48 h, followed by treatment with CldU for 20 min and IdU plus DMSO (con) or 1 μM CPT for 20 min. Data from 1 repeat. B) Replication fork speeds in HCT116 cells (CldU and IdU label) after HMBA treatment +/- HEXIM1 siRNA. n=3. C) First label (CldU) speeds in U2OS cells treated with non-targeting (nonT) or HEXIM1 siRNA for 48 h, followed by treatment with CldU for 20 min and IdU plus DMSO (con), 10 μM CPT or 200 μM HU for 20 min. n=2. D) First label (CldU) speeds in U2OS cells treated with non-targeting (nonT) or LARP7 siRNA for 48 h, followed by treatment with CldU for 20 min and IdU plus DMSO (con), 10 μM CPT or 200 μM HU for 20 min. n=2-3.
Scatter graphs show aggregate of independent repeat with median (line). Column graphs show mean ± SEM. ANOVA or Kruskal-Wallis with multiple comparisons test, * $p \leq 0.05$; **** $p \leq 0.0001$; ns: not significant.

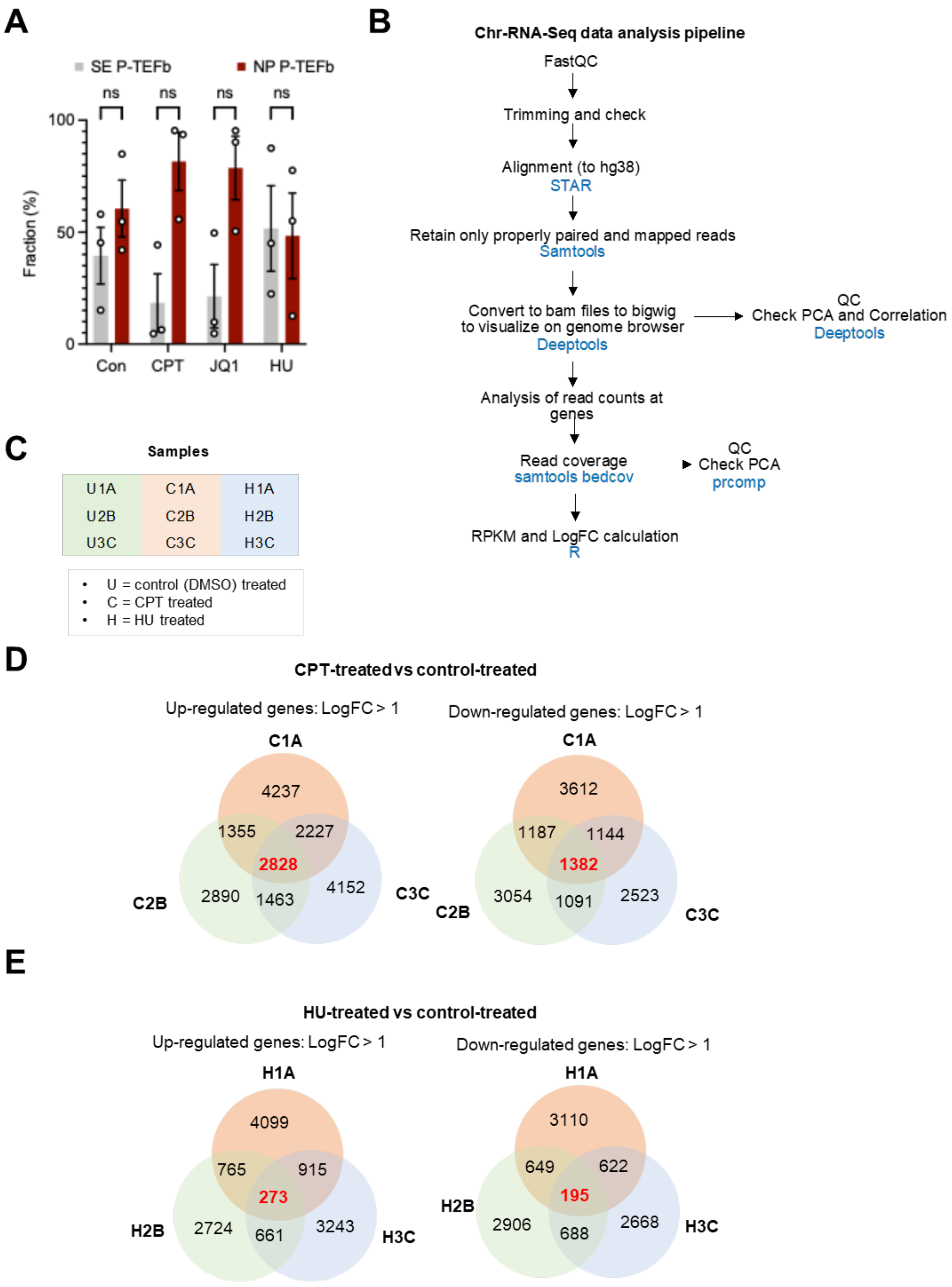

**Figure S2. Impact of CPT and HU on the relocation of P-TEFb and nascent RNA synthesis.**
A) Quantification of relative CDK9 band intensity in U2OS cells after 2 h treatment with 10 µM CPT, 200 µM HU or 1 µM JQ1. n=3. B) Workflow for computational analysis of chromatin RNA-seq. C) Schematic of treatment conditions used for chromatin RNA-seq. D) Venn diagrams of up-regulated and down-regulated in CPT- versus control-treated cells. E) Venn diagrams of up-regulated and down-regulated in HU- versus control-treated cells.
Column graphs show mean ± SEM. ANOVA with multiple comparisons test, ns: not significant.

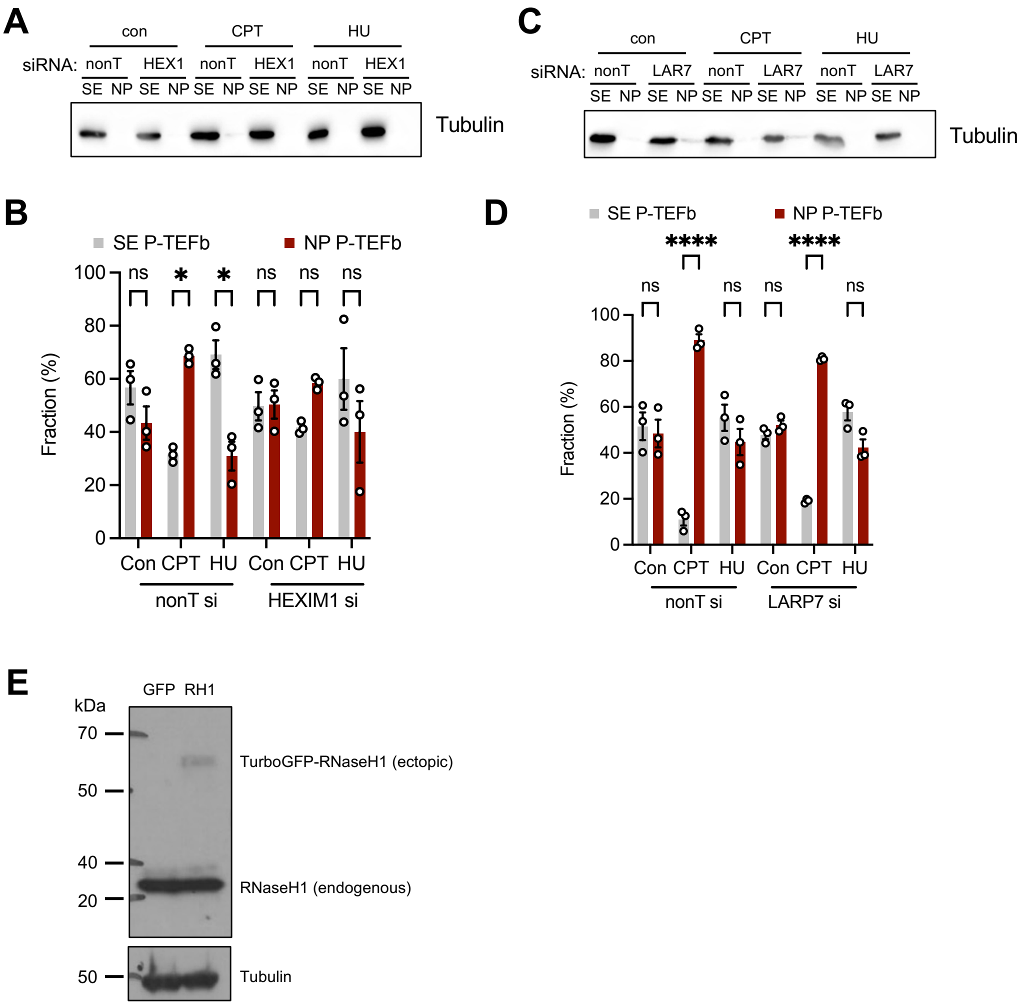

**Figure S3. Impact of HEXIM1 or LARP7 depletion on P-TEFb relocation to chromatin, and ectopic RNase H1 overexpression.**

A) Protein levels of Tubulin (loading control) in cytoplasmic extract (CE) and nuclear pellet (NP) fractions after 2 h treatment with 10 μM CPT or 200 μM HU +/- HEXIM1 siRNA. B) Quantification of relative CDK9 band intensity in U2OS cells treated with non-targeting (nonT) or HEXIM1 siRNA for 48 h, followed by treatment with 10 μM CPT or 200 μM HU (2 h). n=3. C) Protein levels of Tubulin (loading control) in cytoplasmic extract (CE) and nuclear pellet (NP) fractions after 2 h treatment with 10 μM CPT or 200 μM HU +/- LARP7 siRNA. D) Quantification of relative CDK9 band intensity in U2OS cells treated with non-targeting (nonT) or LARP7 siRNA for 48 h, followed by treatment with 10 μM CPT or 200 μM HU (2 h). n=3. E) Protein levels of TurboGFP-RNaseH1, RNaseH1 and Tubulin (control) in U2OS cells 24 h after control (GFP) or TurboGFP-RNaseH1 (RH1) plasmid transfection.

Column graphs show mean ± SEM. ANOVA with multiple comparisons test, * $p \leq 0.05$; **** $p \leq 0.0001$; ns: not significant. Selected p values are shown for clarity.

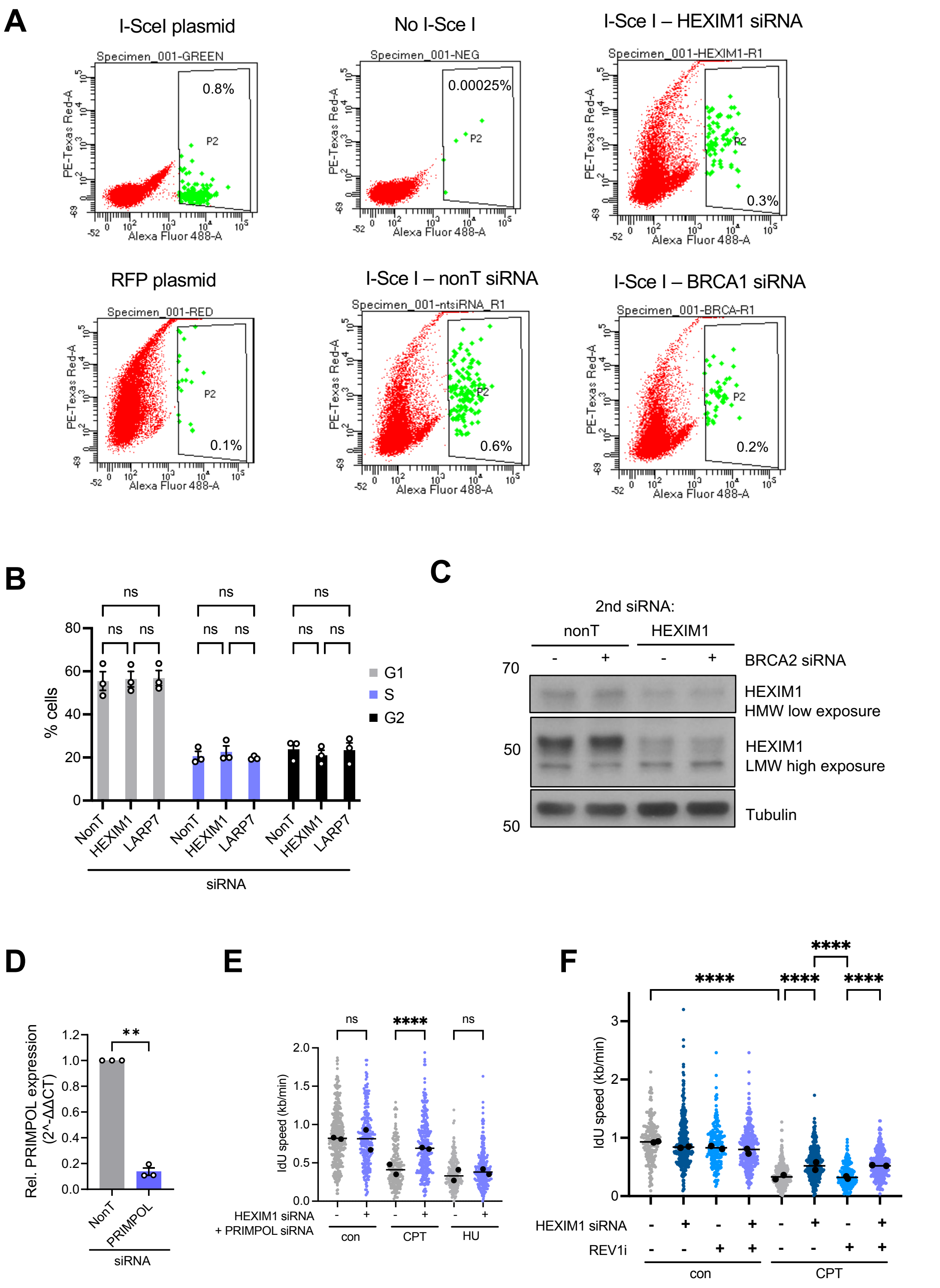

**Figure S4. Role of HEXIM1 in homologous recombination repair and fork reversal.** A) Flow cytometry gating strategy for U2OS-DR-GFP reporter assay, showing representative plots for positive controls (I-Sce I or RFP plasmids), negative control (no I-SceI transfection) and 72 h I-Sce I transfection with non-targeting (nonT), HEXIM1 or BRCA1 siRNA (96 h). B) Quantification of cell cycle distribution in U2OS cells after 72 h transfection with non-targeting (nonT), HEXIM1 or LARP7 siRNA, based on measuring DNA content using propidium iodide staining and flow cytometry. n=3. C) Protein levels of HEXIM1 and Tubulin (loading control) in U2OS cells in presence of BRCA2 (+) or nonT (-) siRNA +/- HEXIM1 siRNA. HMW: HEXIM1 high molecular weight band, LMW: HEXIM1 low molecular weight band. D) RT-qPCR analysis of *PRIMPOL* expression in U2OS cells after 48 h treatment with non-targeting (nonT) control or PRIMPOL siRNA. n=3. E) Replication fork speeds (IdU label) after CPT treatment +/- HEXIM1 and PRIMPOL siRNA. Data from 2 repeats. F) Replication fork speeds (IdU label) after CPT treatment +/- HEXIM1 siRNA and REV1 inhibitor (REV1i). Data from 2 repeats. Scatter graphs show aggregates and medians (black points) of independent repeats with overall median (line). Column graphs show mean ± SEM. Paired t-test, ANOVA or Kruskal-Wallis with multiple comparisons test, ** $p \leq 0.01$; **** $p \leq 0.0001$; ns: not significant. Selected p values are shown for clarity.

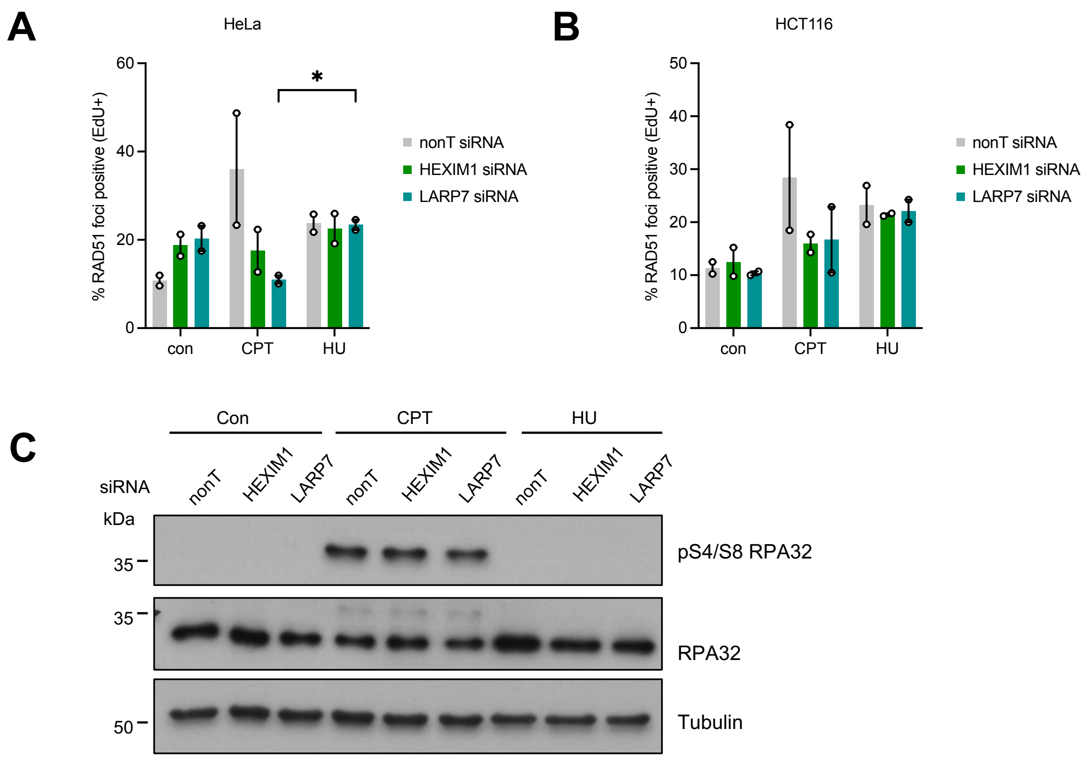

**Figure S5: Impact of HEXIM1 and LARP7 on RAD51 recruitment and DNA end resection.**

A) Percentages of EdU-positive HeLa cells with RAD51 foci after 2 h treatment with 10 μM CPT or 200 μM HU +/- HEXIM1 or LARP7 siRNA. n=2.

B) Percentages of EdU-positive HCT116 cells with RAD51 foci after 2 h treatment with 10 μM CPT or 200 μM HU +/- HEXIM1 or LARP7 siRNA. n=2.

C) Protein levels of phospho-serine 4/8 RPA32, RPA32 (loading control), and Tubulin (control) after 2 h treatment with 10 μM CPT or 200 μM HU +/- HEXIM1 or LARP7 siRNA. The blot shows a different experiment from that in Figure 5I.

Column graphs show mean ± SEM. ANOVA with multiple comparisons test, * $p \leq 0.05$; ** $p \leq 0.01$; *** $p \leq 0.001$; **** $p \leq 0.0001$; ns: not significant.

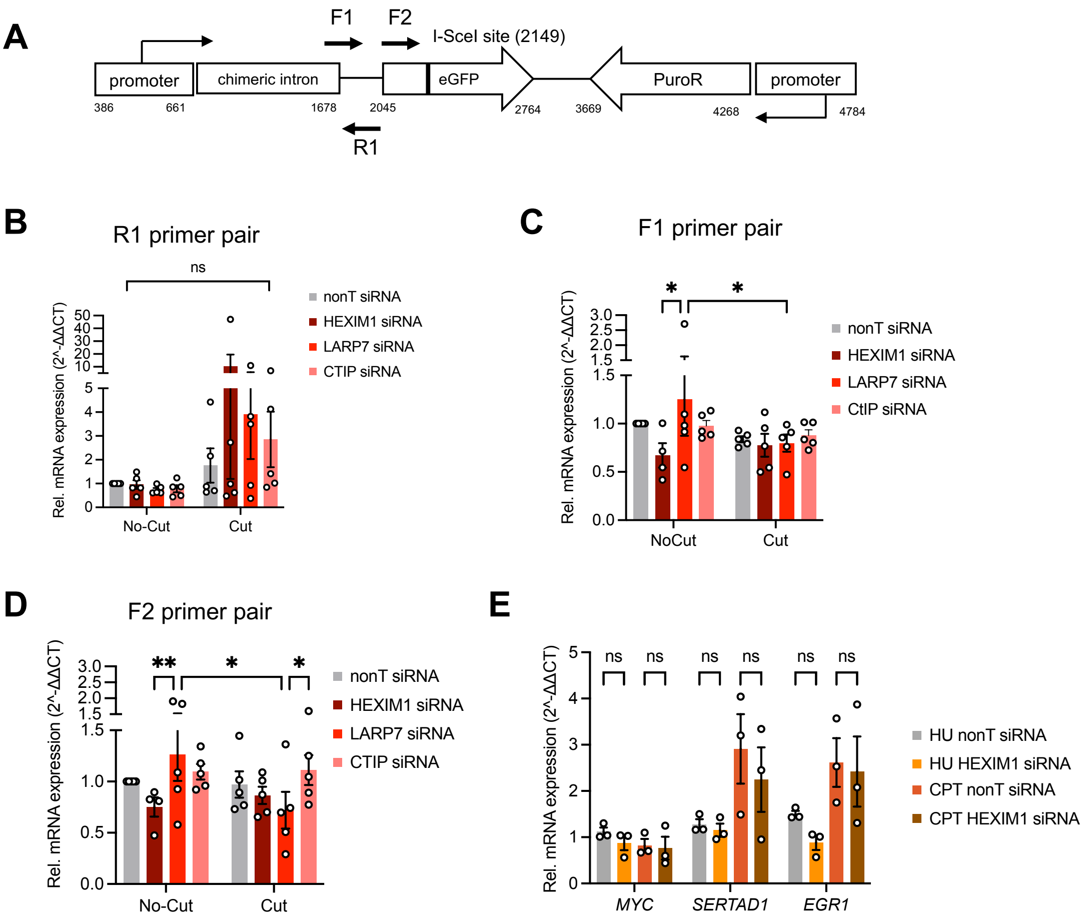

**Figure S6. Impact of HEXIM1 and LARP7 depletion on transcription in the HR reporter construct.**

A) Schematic of a section of the DR-GFP reporter construct with locations of eGFP and PuroR genes and their promoters near the I-SceI restriction site. Arrows show location and direction of transcripts amplified with the F1, F2 or R1 primer pairs. B) RT-qPCR analysis of transcript expression upstream of the eGFP gene in U2OS-DR-GFP cells treated with non-targeting (nonT), HEXIM1, LARP7 or CtIP siRNA for 24 h, followed by transfection (cut) or no transfection (no-cut) with RFP and I-SceI expression constructs for 72 h. n=4-5. C) RT-qPCR analysis of transcript expression at the eGFP gene in U2OS-DR-GFP cells treated as in D. n=4-5. D) RT-qPCR analysis of antisense transcript expression upstream of the eGFP gene in U2OS-DR-GFP cells treated as in D. n=5. E) RT-qPCR analysis of *MYC*, *SERTAD1* and *EGR1* expression after 2 h treatment with 200 200 μM HU or 10 μM CPT +/- HEXIM1 siRNA. n=3.

Column graphs show mean ± SEM. ANOVA with multiple comparisons test, * $p \leq 0.05$; ** $p \leq 0.01$; ns: not significant.

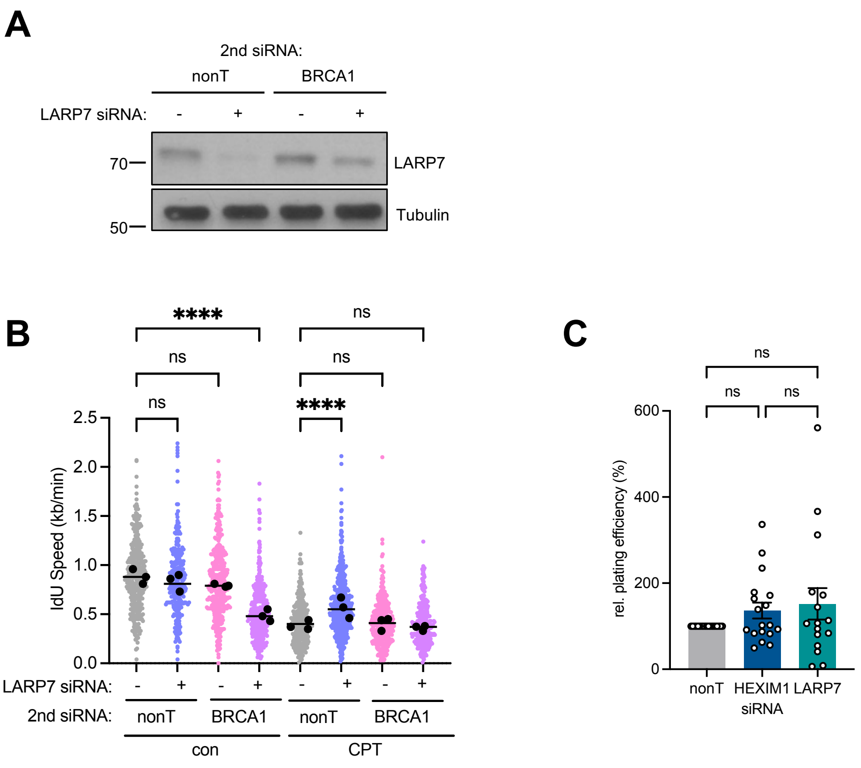

**Figure S7. Relationship of LARP7 with BRCA1 and impact on colony formation**

A) Protein levels of LARP7 and Tubulin (loading control) in U2OS cells 48 h after LARP7 and/or BRCA1 siRNA transfection. B) Second label (IdU) speeds in U2OS cells after CPT treatment +/- BRCA1 and/or LARP7 siRNA. Data from 3 repeats. C) Percentage plating efficiency of U2OS cells treated with HEXIM1 or LARP7 siRNA in colony assays, relative to non-targeting (nonT) control siRNA. n=18 (HEXIM1 siRNA); n = 16 (LARP7 siRNA).

Scatter blots show aggregates and medians (black points) of independent repeats with overall median (line). Column graphs show mean ± SEM. ANOVA or Kruskal-Wallis with multiple comparisons test, **** $p \leq 0.0001$; ns: not significant.

**Supplementary Table S2. Sequences of PCR primers**

| Name | Sequence (5' → 3') | Annealing temp (°C) |
|---|---|---|
| **EGR1 Forward** | TGCAGATCTCTGACCCGTTC | 60 |
| **EGR1 Reverse** | CAGGAAAAGACTCTGCGGTCA | |
| **SERTAD1 Forward** | GAGGACAGCCAACAAGCGAT | 60 |
| **SERTAD1 Reverse** | TGCTCAGCATCTTGCTCACTA | |
| **MYC Forward** | CAGCGACTCTGAGGAGGAAC | 60 |
| **MYC Reverse** | GCTGCGTAGTTGTGCTGATG | |
| **PRIMPOL Forward** | AAAAGCAACGTGGGGCATTC | 55 |
| **PRIMPOL Reverse** | GGTGGTTCTTCTGGCTTGGA | |
| **DR-GFP (F1) Forward** | GTGATCAGGCAGAGCAGGAA | 58 |
| **DR-GFP (F1) Reverse** | ACTTGTGGCCGTTTACGTCG | |
| **DR-GFP (F2) Forward** | TGAGCAAGGGCGAGGAGCTGTT | 60 |
| **DR-GFP (F2) Reverse** | ACACGCTGAACTTGTGGCCGTTTA | |
| **DR-GFP (R1) Forward** | GTCCCCTTCTCCCTCTCCAG | 58 |
| **DR-GFP (R1) Reverse** | TGAACATGGTTAGCAGAGGCT | |
| **BRCA1 Forward** | GCCAAGGCAAGATCTAGAGG | 57 |
| **BRCA1 Reverse** | GTTGCCAACACGAGCTGA | |
| **BRCA2 Forward** | CACCTCTGGAGCGGACTTATT | 57 |
| **BRCA2 Reverse** | GCTTTGTTGCAGCGTGTCTT | |
| **RPLP0 Forward (control)** | CAGATTGGCTACCCAACTGTT | 60 |
| **RPLP0 Reverse (control)** | GGAAGGTGTAATCCGTCTCCAC | |
| **ACTB Forward (control)** | TTGCGTTACACCCTTTCTTG | 60 |
| **ACTB Reverse (control)** | CACCTTCACCGTTCCAGTTT | |
